# Direct carbon monoxide fixation via the bacterial and archaeal Wood–Ljungdahl pathways

**DOI:** 10.1101/2025.10.29.685450

**Authors:** Yuto Fukuyama, Sanae Sakai, Yoko Chiba, Shigeru Shimamura, Satoshi Hiraoka, Tomoyuki Wakashima, Takuro Nunoura, Ken Takai

**Affiliations:** Institute for Extra-cutting-edge Science and Technology Avant-garde Research (X-star), Japan Agency for Marine-Earth Science and Technology (JAMSTEC), 2–15 Natsushima-cho, Yokosuka, Kanagawa, 237–0061, Japan; Research Center for Bioscience and Nanoscience (CeBN), Research Institute for Marine Resources Utilization (MRU), Japan Agency for Marine-Earth Science and Technology (JAMSTEC), 2–15 Natsushima-cho, Yokosuka, Kanagawa, 237–0061, Japan; RIKEN Center for Sustainable Resource Science, 2-1 Hirosawa, Wako, Saitama, 351-0198, Japan; Graduate School of Science and Technology, University of Tsukuba, 1-1-1 Tennodai, Tsukuba, Ibaraki, 305-8577, Japan

## Abstract

The Wood–Ljungdahl (WL) pathway, which is widely distributed in both archaea and bacteria, is an ancient carbon fixation pathway from CO_2_. CO_2_ fixation proceeds via two branches of pathway: progressive reduction to the methyl group in the methyl branch and one-step reduction to CO in the carbonyl branch. In the final step of the pathway, the methyl group, CO, and CoA are combined into a carbon monoxide dehydrogenase (CODH)/acetyl-CoA synthase (ACS) complex to form acetyl-CoA. Here, we show direct CO fixation to the carbonyl group of acetyl-CoA in both archaeal and bacterial WL pathways under hydrogenogenic growth conditions using ^13^C tracer-based metabolomics. A combination of metabolomics and proteomics suggested that the hydrogenotrophically grown *Thermodesulfatator indicus* and *Archaeoglobus* sp. strain MCR cells, directly fixed CO using free-form ACS in the carbonyl branch with relatively low CO availability. In contrast, carboxydotrophically grown *Archaeoglobus* cells utilize the CODH/ACS complex for CO_2_ fixation rather than CO fixation. Direct CO fixation by free-form ACS is more advantageous for conserving reduced ferredoxin compared with the thermodynamically challenged CO_2_ reduction by CODH. These findings provide further insight into the origin and evolution of the most ancient inorganic carbon fixation pathway and geochemical cycles on early Earth.

## Introduction

The Wood–Ljungdahl (WL) pathway is known to be one of the most ancient inorganic carbon fixation pathways distributed in archaea and bacteria^1, 2, 3, 4, 5^. Noncyclic CO_2_ fixation pathway is shared by both archaeal and bacterial WL pathways where CO_2_ is reduced to a methyl group (CH_3_-) progressively in the methyl branch and to CO in the carbonyl branch. However, archaeal and bacterial pathways differ in the enzymes and cofactor carriers in the methyl branch^6^. Finally, acetyl-CoA is formed by a complex of anaerobic CO dehydrogenase/acetyl-CoA synthase (CODH/ACS)^7, 8, 9^. CODH reduces CO_2_ to CO using reduced ferredoxin, and ACS synthesizes acetyl-CoA from CO, a methyl group, and CoA^10^.

If the WL pathway is a primordial CO_2_ fixation system, several critical questions arise. Recently, a three-dimensional structure-based phylogenetic analysis of the hybrid cluster protein (HCP)/CODH superfamily suggested that the primordial enzymes of the HCP/CODH superfamily lack CODH activity^11^. This finding contradicts the hypothesis that the WL pathway is the primordial CO_2_ fixation pathway on Earth. In addition, the largest thermodynamic barrier in the WL pathway is present in the reactions with the CODH/ACS complex: the reduction of CO_2_ to CO using the reduced ferredoxin has a standard redox potential of –524 mV to –558 mV^12^. In modern organisms fixing carbon by the WL pathway, to overcome the barriers, flavin-based electron bifurcation (FBEB) enzymes generally facilitate ferredoxin turnover^13, 14, 15^. As a result, H_2_ oxidation, with a standard redox potential of –414 mV, supplies reduced ferredoxin for carbon fixation via the WL pathway^13^. However, phylogenetic analysis of bifurcating enzymes suggests that the complex and mature machinery are not used for energy metabolism for the primordial CO_2_ fixation^16^.

Anaerobic archaea and bacteria that can use CO as an energy source with relatively high CO availability (carboxydotrophs) have been reported^17^. They possess phylogenetically distinct CODHs from widely distributed CODHs that are responsible for carbon fixation^18, 19, 20^. Because of the notably low redox potential of CO, CO oxidation can couple with the reduction of most electron acceptors, including ferredoxin^21^ using other CODHs responsible for energy conservation^17, 22^. In contrast, CO can be used as the carbon source as an intermediate product in the WL pathway. However, knowledge of CO utilization as the carbon source or both energy and carbon sources in carboxydotrophs remains in its infancy. In this study, to elucidate the metabolic function of the WL pathway in carboxydotrophic and non-carboxydotrophic organisms, we performed ^13^C tracer-based metabolomics using microfluidic capillary electrophoresis-mass spectrometry (CE–MS) on a non-carboxydotrophic, hydrogenotrophic, and sulfate-reducing chemolithoautotrophic bacterium, *Thermodesulfatator indicus*, and a facultative carboxydotrophic and obligatory chemolithoautotrophic archaeon, *Archaeoglobus* sp. strain MCR. This study demonstrates a novel carbon fixation mechanism that incorporates CO directly, rather than via CO_2_ reduction, in both archaea and bacteria under hydrogenotrophic growth conditions. These findings provide a key clue to inorganic carbon fixation via the primordial WL pathway on early Earth.

## Results

### Characteristics of energy and carbon metabolisms

*Thermodesulfatator indicus* and *Archaeoglobus* sp. strain MCR were used as representative hydrogenotrophic bacteria and archaea, respectively, which potentially grow with via WL pathway for inorganic carbon fixation based on their growth characteristics and genome sequences. *T. indicus* is an anaerobic and hydrogenotrophic sulfate-reducer that grows chemolithoautotrophically or chemolithomixotrophically using the WL pathway for inorganic carbon fixation and an incomplete tricarboxylic acid (TCA) cycle for essential intermediates synthesis^23, 24^. In the anaerobic and sulfate-reducing archaeal genus *Archaeoglobus*, some species and strains grow chemolithoautotrophically under hydrogenotrophic conditions^25, 26^. *Archaeoglobus fulgidus* also exhibits carboxydotrophic growth with acetate production^27, 28^. The *Archaeoglobus* sp. strain MCR is a novel isolate from a deep-sea hydrothermal vent ecosystem at the Mid-Cayman Ridge Spreading Center. This strain was isolated as a hydrogenotrophic sulfate reducer and grows carboxydotrophically under an atmosphere of up to 65% CO at 300 kPa. Under a headspace gas of N_2_:CO:CO_2_ (30:50:20) at 300 kPa, the archaeon oxidized CO coupling with H_2_ production and little acetate production (Fig. S1), whereas no H_2_ production was detected under carboxydotrophic growth conditions with acetate production in *A. fulgidus*^27^ (Table S1). Furthermore, the MCR strain utilized CO as the sole carbon source and electron donor coupled with sulfate reduction. The requirement for CO_2_ under carboxydotrophic conditions in the absence of sulfate was shared by *A. fulgidus* (Table S1).

### Phylogeny and gene organization of CODHs

To elucidate CO-related metabolic functions in *T. indicus* and *Archaeoglobus* sp. strain MCR, CODH genes were examined from their genome sequences. Among the CODHs, a type of CODH that is phylogenetically distinct from the CODH involved in the CODH/ACS complex^18, 19^, forms a complex with energy-converting hydrogenase (ECH)^29, 30^. The CODH/ECH complex couples CO oxidation to H_2_ production, generating an electrochemical proton gradient across the cytoplasmic membrane for ATP synthesis^30,31^. Thus, the complex is known to function in carboxydotrophic bacteria and archaea that utilize CO as an energy source under the growth conditions with relatively high CO partial pressures^17^. Although CODH genes are phylogenetically diverse, they frequently form a gene cluster with associated functional genes, such as the CODH−ACS and CODH−ECH gene clusters^18, 19, 20^. Therefore, to estimate the metabolic functions of the proteins encoded in the orthologues of the CODH gene, a combination of phylogenetic analysis and information on the gene organization of the gene cluster, including the CODH gene, is necessary^18, 19^.

We retrieved five (ALIMCR_05470, ALIMCR_12620, ALIMCR_12740, ALIMCR_13930, and ALIMCR_16380) and two (Thein_1422 [WP_013908030.1] and Thein_2190 [WP_013908774.1]) CODH candidates from *Archaeoglobus* sp. strain MCR and *T. indicus* genomes, respectively, using a CODH database^20^ and sequence similarity-based search. The candidates from *Archaeoglobus* sp. strain MCR were spread through phylogenetic clades (Clades A, C, D, and E), whereas the two from *T. indicus* were classified into Clade E (Fig. S2). Active sites conserved among the reference CODHs^19^ are absent in ALIMCR_12620, ALIMCR_12740, and ALIMCR_16380 (Fig. S3). The gene organization around Thein_1422, which harbors conserved active sites, was distinct from known functional CODHs^19^. In contrast, ALIMCR_05470 and Thein_2190 were found in well-known archaeal and bacterial types of the CODH–ACS gene cluster (ALIMCR_05430–ALIMCR_05520 and Thein_2175–Thein_2192, respectively) (Table S2). In addition, ALIMCR_13930 was located in the gene cluster of membrane-bound energy-converting hydrogenase (Mbh–type ECH) (Table S3) (ALIMCR_13910– ALIMCR_14030). Although a transcriptional regulator gene was not identified, the gene organization was similar to that of the genomes of hydrogenogenic and carboxydotrophic strains of the archaeal genus *Thermococcus*^32, 33^ (Fig. S4). Thus, the CODH/Mbh–type ECH complex in *Archaeoglobus* sp. strain MCR likely plays essential roles in their hydrogenogenic carboxydotrophy. In contrast, *A. fulgidus* harbors three CODH genes, including two archaeal-type genes, but lacks the CODH gene for the CODH/Mbh–type ECH complex^18, 19^. One of the archaeal-type CODH genes functions as a CODH/ACS complex, whereas the other oxidizes CO coupled with the reduction of oxidized ferredoxin^28^. Thus, the carboxydotrophy-related system in *A. fulgidus* was distinct from that of strain MCR.

### CO impact on protein expression in strain MCR

To identify the proteins involved in energy and central carbon metabolism, especially CODHs, under hydrogenotrophic and carboxydotrophic growth conditions, we performed comparative semi-quantitative shotgun proteomics on *Archaeoglobus* sp. strain MCR (Fig. S5). With the exception of succinate dehydrogenase, proteins involved in an incomplete TCA cycle were detected under all growth conditions. Of these, an abundance of enzymes, such as pyruvate ferredoxin oxidoreductase (PorABCD; ALIMCR_12950–12980), pyruvate carboxylase (PycAB; ALIMCR_04860 and ALIMCR_06820), and malate dehydrogenase (MDH; ALIMCR_06530) was greater under carboxydotrophic conditions than hydrogenotrophic conditions. Proteins involved in the archaeal WL pathway, including those encoded by genes in the CODH–ACS gene cluster (ALIMCR_05440–05510), were also maintained under all growth conditions. In contrast, proteins involved in the CODH/Mbh–type ECH complex (ALIMCR_13910– 14030) were limited to cells grown in the absence of CO, whereas they were highly expressed in cells grown under carboxydotrophic conditions. These results revealed that the CODH/Mbh–type ECH complex was regulated in a CO-dependent manner. Proteins involved in sulfate respiration and hydrogenase activity were relatively suppressed under carboxydotrophic conditions. Membrane-bound redox complex proteins (DsrMKJOP; ALIMCR_1069-1074) were depleted with sulfate unavailability.

### Direct CO fixation via the bacterial WL pathway

We performed ^13^C tracer-based metabolomics using CE–MS that targeted proteinogenic amino acids for isotopologue and isotopomer analyses^34^. Recent ^13^C tracer-based metabolomics of *T. indicus* has shown that CO_2_ is fixed via the bacterial WL pathway^24^. The generated acetyl-CoA is incorporated into the incomplete and bifurcated TCA cycle to synthesize amino acids^24^. Genomic information suggests that alanine (Ala), aspartate (Asp), and glutamate (Glu) are synthesized from pyruvate, oxaloacetate, and 2-oxoglutarate, respectively, and the results of tracer-based metabolomics are consistent with the expected pathways^24^. To obtain further insights into the operation of the WL pathway and CO fixation machinery, we conducted tracer-based metabolomics in this study.

All isotopologues of Ala, Asp, and Glu were detected in *T. indicus* cells grown under a headspace gas of H_2_:^12^CO_2_:^13^CO_2_ (80:10:10) (Fig. 1A and Table 1). From the isotopomer analysis of the amino acids containing one ^13^C-labeled carbon, the relative abundance of ^13^C-labeled carbon in each fragment was greater than that of natural ^13^C abundance (1.1%), and their proportions were similar (Figs. S6-8). For example, in the case of Asp, the relative abundance of ^13^C-labeled carbon at all positions was greater than that of the negative control with equal abundance (Fig. 1), suggesting that ^13^CO_2_ was incorporated into any positions of Asp. A limitation of isotopomer analysis is its inability to distinguish between ^13^C-labeled carbon at positions 3 and 4. These equal distributions of ^13^C can be explained by the fixation of two molecules of CO_2_ via the methyl and carbonyl branches in the bacterial WL pathway, followed by oxaloacetate synthesis from acetyl-CoA via pyruvate^34^. The CO_2_ molecules that are fixed in the carboxylation of acetyl-CoA, carbonyl branch, methyl branch, and carboxylation of pyruvate are arranged from the first to the fourth position (Fig. 1B). Accordingly, the isotopologue and isotopomer patterns were consistent with chemolithoautotrophic inorganic carbon fixation via the WL pathway in *T. indicus*.

**Fig. 1.**
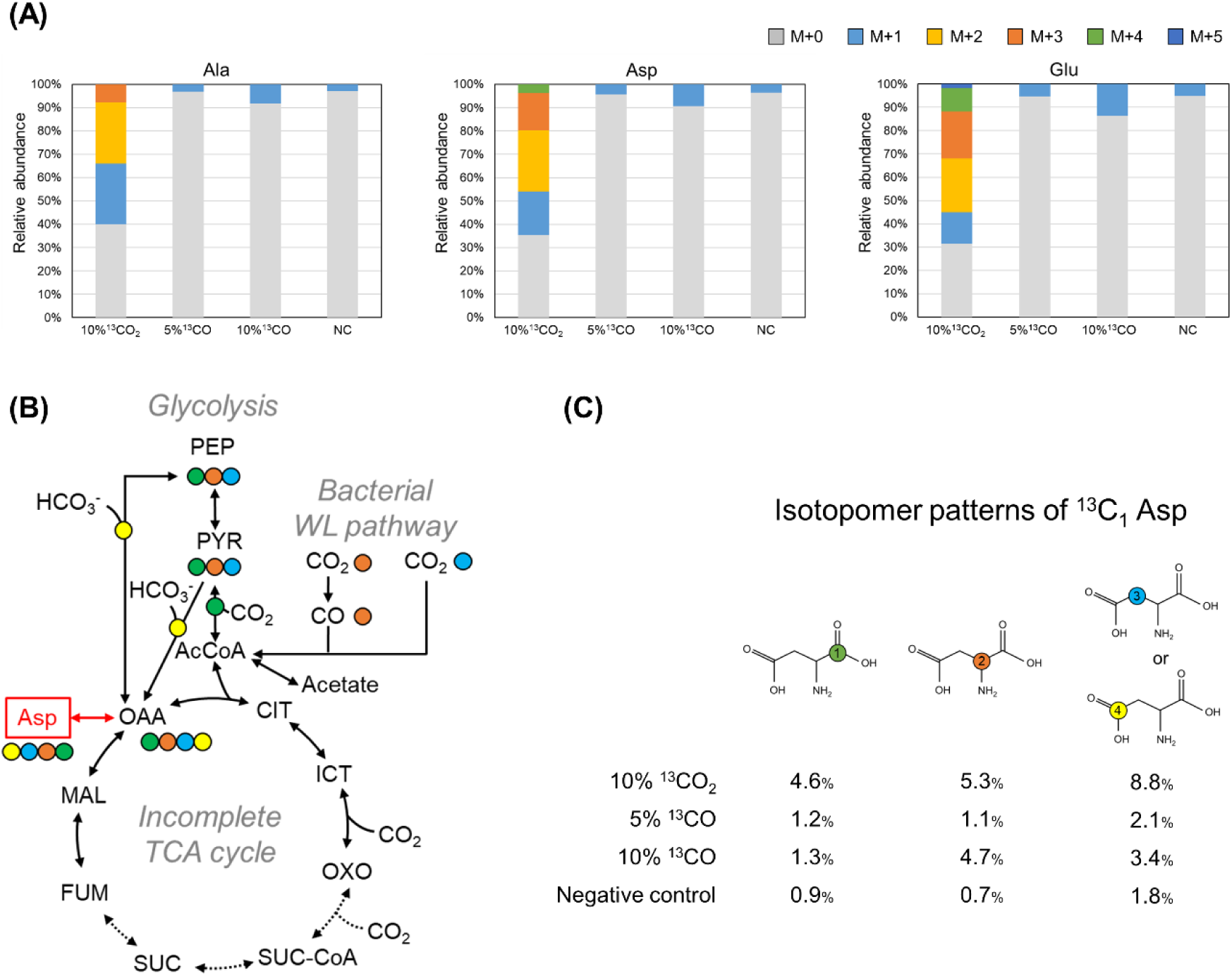
**Mass isotopomer distributions from Ala, Asp, and Glu (A), proposed carbon flow from metabolome analysis (B), and relative abundance of isotopomer patterns in Asp containing one ^13^C-labeled carbon (C) from *Thermodesulfatator indicus* grown under chemolithoautotrophic conditions** The mass fractions for M + 0, M + 1, M + 2, M + 3, M+4, and M+5 represent fragments containing 0–5 ^13^C-labeled carbons. Each number in the circle indicates the carbon position of Asp. PEP, phosphoenolpyruvate; PYR, pyruvate; AcCoA, acetyl-CoA; CIT, citrate; ICT, isocitrate; OXO, 2-oxoglutarate; SUC-CoA, succinyl-CoA; SUC, succinate; FUM, fumarate; MAL, malate; OAA, oxaloacetate; Ala, alanine; Asp, aspartate; Glu, glutamate

**Table 1.** **The relative abundance of mass fractions of Ala, Asp, and Glu from *T. indicus***

| Amino acid | Natural abundance of isotopes | 10% $^{13}\text{CO}_2$ | 5% $^{13}\text{CO}$ | 10% $^{13}\text{CO}$ | Negative control |
| --- | --- | --- | --- | --- | --- |
|  | abundance (%) | abundance (%) | abundance (%) | abundance (%) | abundance (%) |
| $^{13}\text{C}_0$ Ala | 96.7 | 40.0 | 96.9 | 91.9 | 97.1 |
| $^{13}\text{C}_1$ Ala | 3.2 | 26.0 | 3.1 | 8.1 | 2.9 |
| $^{13}\text{C}_2$ Ala | 0.0 | 26.3 | N.D. | N.D. | N.D. |
| $^{13}\text{C}_3$ Ala | 0.0 | 7.7 | N.D. | N.D. | N.D. |
| $^{13}\text{C}_0$ Asp | 95.7 | 35.3 | 95.6 | 90.6 | 96.6 |
| $^{13}\text{C}_1$ Asp | 4.3 | 18.7 | 4.4 | 9.4 | 3.4 |
| $^{13}\text{C}_2$ Asp | 0.1 | 26.3 | N.D. | N.D. | N.D. |
| $^{13}\text{C}_3$ Asp | 0.0 | 15.9 | N.D. | N.D. | N.D. |
| $^{13}\text{C}_4$ Asp | 0.0 | 3.8 | N.D. | N.D. | N.D. |
| $^{13}\text{C}_0$ Glu | 94.6 | 31.3 | 94.7 | 86.4 | 94.8 |
| $^{13}\text{C}_1$ Glu | 5.3 | 13.6 | 5.3 | 13.6 | 5.2 |
| $^{13}\text{C}_2$ Glu | 0.1 | 23.0 | N.D. | N.D. | N.D. |
| $^{13}\text{C}_3$ Glu | 0.0 | 20.2 | N.D. | N.D. | N.D. |
| $^{13}\text{C}_4$ Glu | 0.0 | 9.9 | N.D. | N.D. | N.D. |
| $^{13}\text{C}_5$ Glu | 0.0 | 1.9 | N.D. | N.D. | N.D. |

N.D., not detected

We further investigated the impact of CO as a carbon source on carbon fixation under headspace gas of H_2_:^12^CO_2_:^13^CO (80:10:10 or 85:10:5). The relative abundance of each amino acid containing one ^13^C-labeled carbon increased in *T. indicus* cells along with ^13^CO concentrations (Fig. 1A and Table 1), implying that CO was directly incorporated as a carbon source. If ^13^CO is oxidized to ^13^CO_2_ and then incorporated via the bacterial WL pathway, the relative abundance of ^13^C at each position should be similar. However, the isotopomer pattern of these amino acids in cells grown with ^13^CO was distinct from that in cells grown with ^13^CO_2_ (Figs. S7-9). For example, in the case of 10% ^13^CO addition, the relative abundance of ^13^C-labeled carbon at position 2 in Asp (4.7%) was much higher than that in the negative control (0.7%), although the relative abundance of ^13^C-labeled carbon at other positions increased by less than two-fold (Fig. 1C). These results suggest that CO is directly incorporated into the bacterial WL pathway as a carbon source.

### CO and CO_2_ fixations via the archaeal WL pathway

Based on genomic and proteomic information, *Archaeoglobus* sp. strain MCR likely fixes CO_2_ via the archaeal WL pathway and operates an incomplete TCA cycle. To verify the CO_2_ fixation route and the ability of CO fixation via the archaeal WL pathway, we performed ^13^C tracer-based metabolomics in *Archaeoglobus* sp. strain MCR growing hydrogenotrophically under a headspace gas of H_2_:^12^CO_2_:^13^CO_2_ (80:15:5, 0.3 MPa) or H_2_:^12^CO_2_:^13^CO (80:15:5, 0.3MPa). Up to three ^13^C-labeled carbons were detected in Ala, Asp, and Glu in cells grown hydrogenotrophically with ^13^CO_2_ (Fig. 2A and Table 2). Based on isotopomer analysis, the relative abundance of ^13^C in each amino acid derived from ^13^CO_2_ was similar (Figs. S9-11), which was similar to that of *T. indicus*. In Asp containing one ^13^C-labeled carbon, the relative abundances at all positions were similar and greater than those of the negative control (Fig. 2C). In addition, up to three ^13^C-labeled carbon atoms derived from ^13^CO_2_ were detected, which were distributed in both main- and side-chains of Glu (Fig. 2A and S11, Table 2). If a complete reductive TCA cycle occurs, 2-oxoglutarate (a precursor of Glu) is synthesized from oxaloacetate (a precursor of Asp) via succinyl-CoA. Thus, the presence of M+4 and M+5 Glu is expected. However, these isotopologues were absent, and a tiny amount of M+3 in Glu (0.1%) was detected. Accordingly, we concluded that the MCR strain operated the via archaeal WL pathway for carbon fixation along with incomplete TCA cycle, which lacks reactions between succinate and 2-oxoglutarate, as observed in *T. indicus*^24^. When ^13^CO (10%, v/v) was added, up to three ^13^C-labeled carbons in Ala and Asp and up to four ^13^C-labeled carbons in Glu were detected (Table 2), whereas the relative abundance of M+2 in Ala and Asp was low (almost negligible). Isotopomer analysis showed that ^13^CO incorporation rates varied among positions, as observed in *T. indicus* (Figs. S9-11). The relative abundance of ^13^C-labeled carbon at position 2 in Asp (41.2%) was greater than that at other positions (Fig. 2C). These results suggested that CO is directly incorporated into the archaeal WL pathway in hydrogenotrophic cells. However, low abundance of M+2 Ala and Asp suggests that a tiny amount of ^13^CO was oxidized to ^13^CO_2_, followed by CO_2_ reduction in both branches of the WL pathway.

**Fig. 2.**
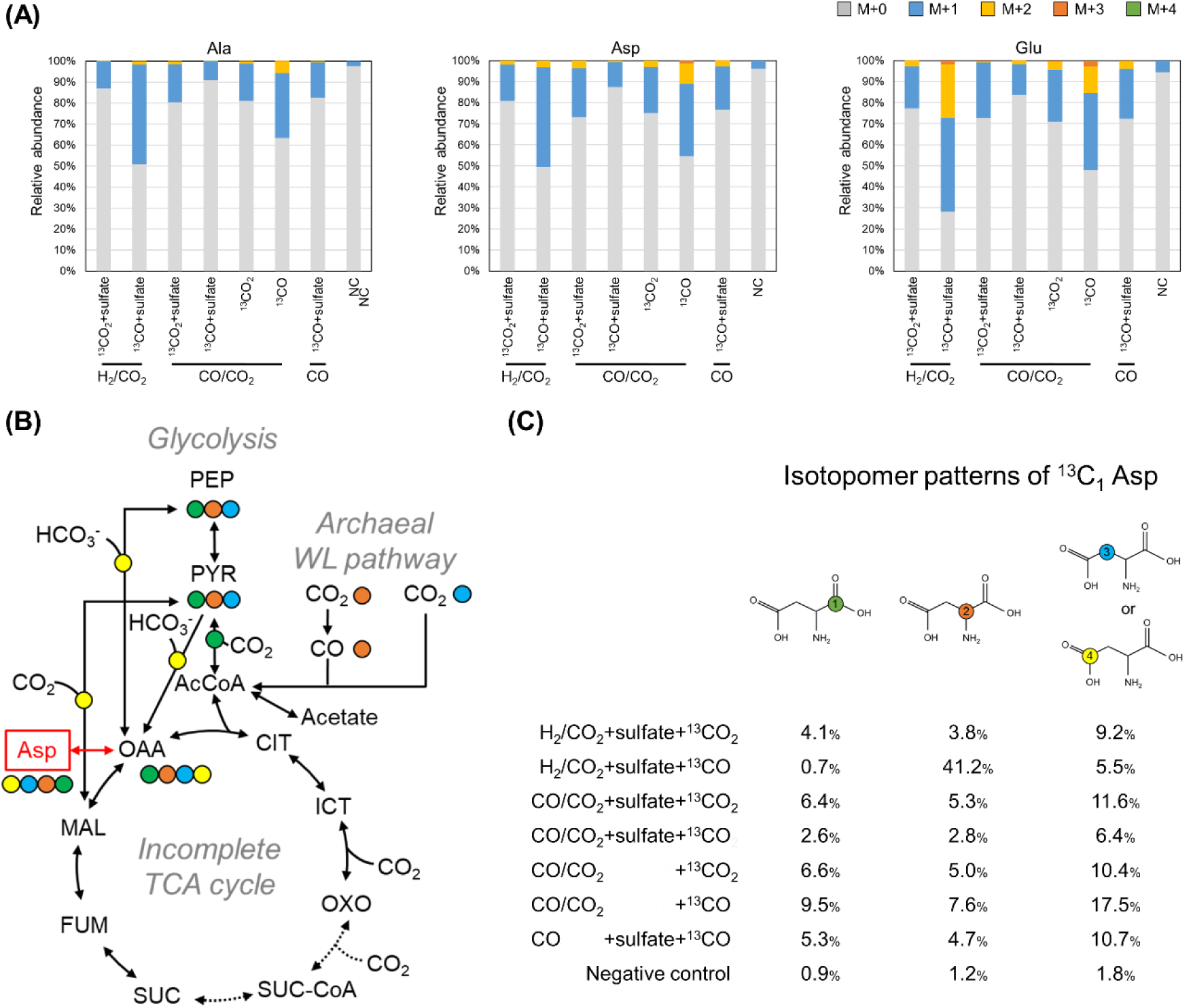
**Mass isotopomer distributions from Ala, Asp, and Glu (A), proposed carbon flow from metabolome analysis (B), and relative abundance of isotopomer patterns in Asp containing one ^13^C-labeled carbon (C) from *Archaeoglobus* sp. strain MCR grown under chemolithoautotrophic or carboxydotrophic conditions** The mass fractions for M + 0, M + 1, M + 2, M + 3, and M+4 represent fragments containing 0–4 ^13^C-labeled carbons, respectively. Each number in the circle indicates the carbon position of Asp. PEP, phosphoenolpyruvate; Pyr, pyruvate; AcCoA, acetyl-CoA; CIT, citrate; ICT, isocitrate; OXO, 2-oxoglutarate; SUC-CoA, succinyl-CoA; SUC, succinate; FUM, fumarate; MAL, malate; OAA, oxaloacetate; Ala, alanine; Asp, aspartate; Glu, glutamate

**Table 2.**
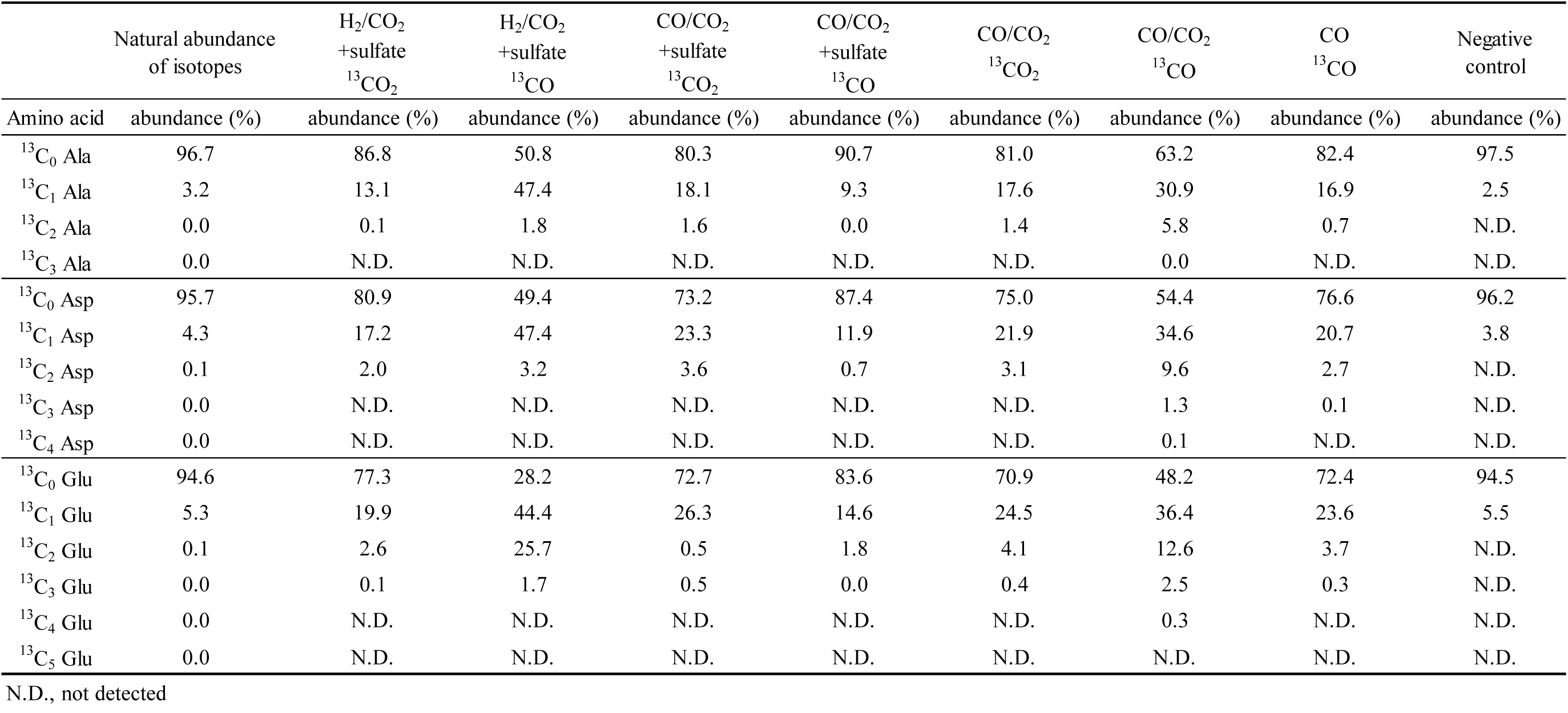
The relative abundance of mass fractions of Ala, Asp, and Glu from Archaeoglobus sp. strain MCR.

To further probe the CO_2_ and CO fixation pathways in *Archaeoglobus* sp. strain MCR under carboxydotrophic growth condition with up to 50% (v/v) CO in the headspace gas (Fig. 2), we conducted isotopologue and isotopomer analyses for cells grown carboxydotrophically under a headspace gas of N_2_:^12^CO:^12^CO_2_:^13^CO_2_ (30:50:15:5) or N_2_:^12^CO:^12^CO_2_:^13^CO (30:45:20:5) with or without sulfate. The results showed that the relative abundance of ^13^C-labeled carbon at each position was similar to each other in Asp in cells grown with ^13^CO_2_ (5%, v/v) addition (Fig. 2C). Thus, we conclude that the metabolic function of the WL pathway for CO_2_ fixation was maintained under the carboxydotrophic conditions using CO as the sole electron donor. In contrast, when ^13^CO (5%, v/v) was added to trace incorporation route under carboxydotrophic conditions with or without sulfate, the isotopomer analysis showed that the abundance of ^13^C-labeld carbon was equally distributed among the any positions in Asp (Fig. 2C), as in the case of ^13^CO_2_ addition under carboxydotrophic conditions. These results suggest that ^13^CO was oxidized to ^13^CO_2_ and then incorporated via the archaeal WL pathway under carboxydotrophic conditions.

## Discussion

### Direct CO fixation by free-form ACS

In this study, ^13^C tracer-based metabolomics unveiled direct CO fixation as a carbon source in both *T. indicus* and *Archaeoglobus* sp. strain MCR under hydrogenotrophic growth conditions. The archaeon incorporated CO directly to form acetyl-CoA via the archaeal WL pathways under hydrogenotrophic growth condition with relatively low CO availability, whereas direct CO fixation was not detected under carboxydotrophic growth conditions (Fig. 2). In the CODH/ACS complex, CO_2_ is reduced to CO by CODH, and the generated CO is then transported through a hydrophobic CO tunnel that connects the active sites of both enzymes: the C-cluster in CODH and the A-cluster in ACS^9, 35^. Accordingly, the CODH/ACS complex in both organisms is unlikely to function in direct CO fixation; thus, the free-form ACS is responsible for direct CO fixation in both archaeal and bacterial WL pathways.

In *T. indicus,* during the hydrogenotrophic growth in a headspace gas with 10% ^13^CO_2_ or 10% ^13^CO, the relative abundance of ^13^C-labeled carbon at position 2 in M+1 Asp was 5.3% and 4.7%, respectively (Fig. 1). Meanwhile, that in *Archaeoglobus* sp. strain MCR cells grown hydrogenotrophically with 5% ^13^CO_2_ or 5% ^13^CO was 3.8% and 41.2%, respectively (Fig. 2). This difference suggests that direct CO fixation in archaea is more functional than in *T. indicus*. In addition, considering the low solubility of CO, these results suggest that direct CO incorporation into the carbonyl group of acetyl-CoA in archaeon was more efficient in terms of quality and affinity than CO_2_ incorporation via the carbonyl branch in the WL pathway, and that direct CO fixation in *Archaeoglobus* sp. strain MCR may significantly contribute to biomass production.

### Carboxydotrophic CO and CO_2_ fixation

*Archaeoglobus* sp. strain MCR regulates CO metabolism depending on CO availability. Proteomic analysis indicated CO-driven expression of proteins encoded in the CODH– ECH gene cluster in the archaeon (Fig. S4) similar to other facultative carboxydotrophs^36, 37, 38^. However, a transcriptional regulator-encoding gene was not found (Fig. S4). Fig. 3 presents an overview of the proposed metabolic pathways related to energy and inorganic carbon fixation in archaeon under carboxydotrophic conditions.

**Fig. 3.**
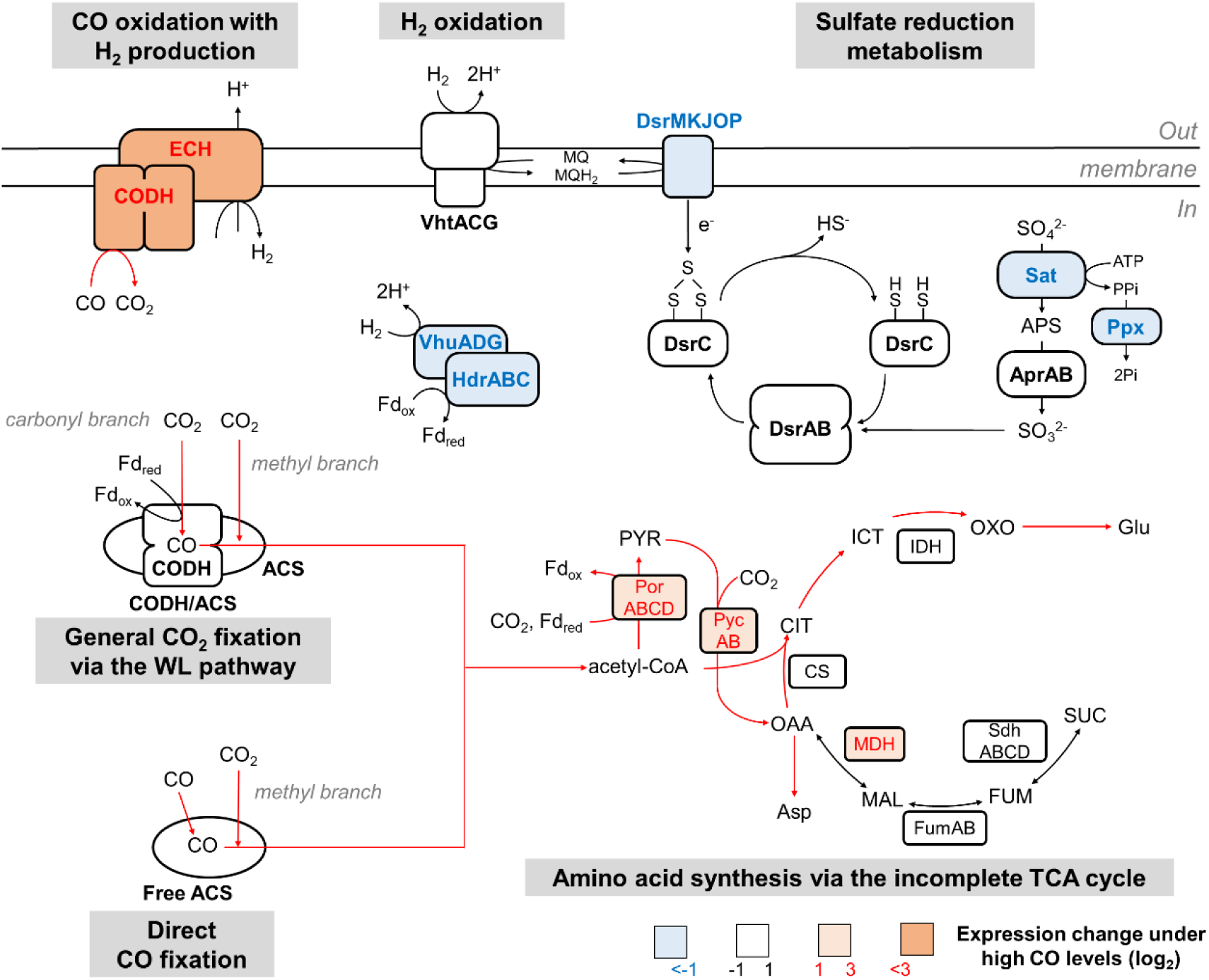
**Overview of the proposed pathways of energy and a part of the central carbon metabolisms in Archaeoglobus sp. strain MCR cells grown hydrogenotrophically or carboxydotrophically** Each log_2_ fold change was calculated between the abundance of protein expression in cells grown under CO/CO_2_ + sulfate and in those grown under H_2_/CO_2_ + sulfate. Red indicates that the log_2_ fold change value is >1. Light blue indicates that the log_2_ fold change value is <-1. The log_2_ fold-change values are listed in the Supporting Information, Fig. S5. CODH, carbon monoxide dehydrogenase; ECH, membrane-bound energy-converting hydrogenase; ACS, acetyl-CoA synthase; VhtACG, methanophenazine hydrogenase; VhuADG, F_420_-non-reducing hydrogenase; HdrABC, soluble heterodisulfide reductase; DsrMKJOP, molybdopterin oxidoreductase; DsrAB and DsrC, dissimilatory sulfite reductase; Sat, sulfate adenylyltransferase; AprAB, adenylylsulfate reductase; Ppx, manganese-dependent inorganic pyrophosphatase; PorABCD, pyruvate-ferredoxin oxidoreductase; PycAB, pyruvate carboxylase; CS, citrate synthase; IDH, isocitrate dehydrogenase; MDH, malate dehydrogenase; FumAB, fumarate hydratase; SdhABCD, succinate dehydrogenase; MQ, reduced menaquinone; MQH_2_, etrahydromethanopterin; Fd, ferredoxin.

The uniqueness of the growth physiology of *Archaeoglobus* sp. strain MCR lies in its ability to utilize CO as the sole carbon source and electron donor when sulfate is available as an electron acceptor, despite the widely accepted view that CO_2_ is necessary for the methyl branch of the WL pathway^6^. In archaeon cells grown carboxydotrophically, CO is oxidized to CO_2_ by the CO-induced highly expressed CODH/Mbh–type ECH complex, which is responsible for coupling CO oxidation with H_2_ production^30, 31, 36^. In contrast to hydrogenotrophic growth, direct CO fixation in carboxydotrophically grown cells was negligible in isotopomer analysis. These results indicates that the abundantly supplied cytosolic CO_2_ by the CODH/Mbh–type ECH complex from CO was incorporated into acetyl-CoA using the CODH/ACS complex, but not directly incorporated with free-form ACS under carboxydotrophic conditions.

### CODHs and ACS regulations and functions

Proteomic analysis suggested that the effects of CO and CO_2_ concentrations were limited to expression of the CODH/ACS complex. This finding is consistent with those of previous studies reporting the purification of the CODH/ACS complex from phylogenetically diverse microbes growing under either hydrogenotrophic or carboxydotrophic conditions^39, 40, 41, 42^. Moreover, the coexistence of monomeric ACS and CODH/ACS complex has also been shown in *Carboxydothermus hydrogenoformans* cells grown carboxydotrophically^43^. Acccordingly, the free-form ACS and CODH/ACS complex would coexist in *T. indicus* and *Archaeoglobus* sp. strain MCR cells grown either hydrogenotrophically or carboxydotrophically.

Only under carboxydotrophic conditions is the archaeon expressing a membrane-bound CODH/Mbh–type ECH complex, and the type of CODHs responsible for energy conservation is known to exhibit notably high CO oxidation activity^17, 44^. In addition, CO resistance acquisition and CODH/ECH complex-associated carboxydotrophy suggest that this complex is responsible for CO detoxification. For example, *Desulfotomaculum nigrificans* strain DSM 574^T^ growth is inhibited under a headspace gas of 20% CO^45^, whereas *D. nigrificans* strain CO-1-SRB, with the CODH/ECH gene cluster, can grow carboxydotrophically under 100% CO with H_2_ production^46, 47^. Accordingly, the most probable scenario for the absence of direct CO fixation signatures under carboxydotrophic conditions in the archaeon is that the cytosolic CO flux under carboxydotrophic conditions was lower because of the CO consumption by the CODH/EDH complex than that under non-carboxydotrophic condition (CO concentration, <20%). Therefore, the ^13^C signature of direct CO fixation was found to be below the threshold (= natural-level ^13^C abundance) used in this analysis.

### Ecological importance of direct CO fixation

Owing to the low redox potential of CO, the advantage of CO oxidation with CODH in energy conservation extends the utilization capability of diverse electron acceptors, including ferredoxin^17, 21, 22^. Moreover, CODH supplies cytosolic CO_2_ for the synthesis of organic compounds via the WL pathway as well as for CO detoxification under carboxydotrophic conditions. In contrast, the advantage of direct CO fixation is that it avoids the thermodynamically difficult step of the WL pathway^6^, endergonic CO_2_ reduction by CODH, which requires reduced ferredoxin. Reduced ferredoxin is also utilized in other essential endogenous reactions, such as pyruvate ferredoxin oxidoreductase (POR), which is involved in gluconeogenesis following the WL pathway. CO uptake by free-form ACS with high CO affinity is a mechanism for reducing the consumption of reduced ferredoxin, thus saving energy under non-carboxydotrophic conditions if CO does not inhibit growth. Thus, simple reactions independent of FBEB are advantageous from the energetic aspect of the WL pathway operation. However, in the case of low cytosolic CO availability, the CODH/ACS complex formation with the high CO affinity of ACS also has an advantage in the endergonic forward reaction of CODH in the WL pathway for CO_2_ fixation to maintain a concentration gradient of CO_2_/CO in the complex. Therefore, the coexistence of free-form ACS and the CODH/ACS complex under both carboxydotrophic and non-carboxydotrophic conditions is an adaptation mechanism for microbial life, with the WL pathway inhabiting relatively CO-enriched hydrothermal and geothermal systems. Moreover, the direct CO fixation machinery implies the origin of the WL pathway. The combination of the CO_2_-fixing methyl branch and the direct CO-fixing carbonyl branch could be one of the most primordial and adapted forms of the WL pathway when early microbial life was originated in the ancient Earth, in which the surface environments (both atmosphere and ocean) were exposed to both high concentrations of CO_2_ as a major inorganic carbon component and CO as a minor, but not negligible, component^48^. Some relic machinery of the primordial WL pathway may have been inherited from modern archaea and bacteria. CODH activity acquisition by an enzyme in the HCP/CODH superfamily may represent the legacy of the primordial WL pathway prior to the development of modern CODHs. Further investigation of genetic and functional diversity and novel variants of the WL pathway will provide important insights into the origin and early evolution of energy and carbon metabolism, as well as the geochemical cycles on ancient Earth.

## Methods

### Strain used in this study

*Archaeoglobus* sp. strain MCR (=DSM 106437) was isolated from a deep-sea hydrothermal vent chimney in the Beebe Vent Field in the Mid-Cayman Spreading Center, Caribbean Sea, and maintained in our laboratory. *T. indicus* strain CIR29812^T^ (=DSM 15286^T^=JCM 11889^T^) was obtained from the Japan Collection of Microorganisms (JCM).

### Proteomic analysis of *Archaeoglobus* sp. strain MCR

*Archaeoglobus* sp. strain MCR cells grown carboxydotrophically with hydrogen production and chemolithoautotrophically with hydrogenotrophic sulfate reduction were used for shotgun proteomic analyses. Proteins were extracted as previously described^49,^_50._

Proteomic analysis was performed using nano flow liquid chromatography–MS (LC–MS), as previously described^49, 50^. Briefly, the digested peptides were analyzed by using the Ultimate 3000 RSLCnano system (Thermo Fisher Scientific) with a reverse-phase Zaplous alpha Pep-C18 column (3 μm, 120 Å, 0.1 × 150 mm; AMR, Tokyo, Japan) coupled with the Orbitrap Fusion Tribrid mass spectrometer. Data analysis of the detected peptides was performed using Proteome Discoverer version 2.2 software package (Thermo Fisher Scientific). Peptides corresponding to a <1% protein false discovery rate (FDR) were used in the calculations. Relative peptide detection values for each protein were calculated. The calculated relative abundance was normalized to the percentage of each protein for comparison between the four tested conditions. The Sum PEP score for each protein was calculated using Proteome Discoverer version 2.2.

### 13C tracer-based metabolomics

All ^13^C-labeled reagents used in this study were purchased from Cambridge Isotope Laboratories (CIL) (Tewksbury, MA, USA).

*T. indicus* was grown in 3 mL of the modified JCM354 medium^24^ under a gas mixture of 80% H_2_ and 20% CO_2_ (0.2 MPa) in 20 mL test tubes at 70°C, as previously described^24^. Unless indicated, *Archaeoglobus* sp. strain MCR was grown in 5 mL of the basal medium (the composition of the medium sees Supplementary Information) under a gas phase of 80% H_2_ and 20% CO_2_ or 20% CO_2_, 50% CO, and 30%N_2_ (0.3 MPa) in 20 mL test tubes at 75°C. For growth without sulfate, Na_2_SO_4_ (2 g/L), MgSO_4_·7H_2_O, and Fe (NH_4_)_2_(SO_4_)_2_·6H_2_O were omitted from the basal medium.

For ^13^C tracer-based metabolomics of *T. indicus*, the cells were grown in modified JCM354 medium supplemented with ^13^C-labeled substrates. After 7 h of cultivation, 4 mL ^13^CO_2_ was added to the headspace of the culture, and then the H_2_ gas was added to 0.2 MPa (final concentration of ^13^CO_2_, 10% [v/v] of the headspace gas). In the case of ^13^CO addition, 1 or 2 mL ^13^CO was added to the headspace gas of H_2_ (0.16 MPa). CO_2_ gas was then added to a pressure of 0.2 MPa, followed by cultivation. As a result, the final concentration of CO was 5% or 10% (v/v) of the headspace gases. For ^13^C tracer-based metabolomics for *Archaeoglobus* sp. strain MCR, 0.9 mL of ^13^CO_2_ or 1.125 mL of ^13^CO were added to the headspace of the culture (final concentration, 5% [v/v] of the headspace). Cells grown with ^13^C-labeled or non-labeled substrates were harvested at the exponential phase by centrifugation (4°C, 10,000 × g, 10 min), frozen in liquid N_2_, and stored at −80°C until use.

Protein-derived amino acids for ^13^C tracer-based metabolomics were prepared as previously described^34^. Briefly, the cells were processed with hydrolyzation in 12N HCl at 110°C for 16 h, followed by liquid-liquid extraction to purify the protein-derived amino acids. Purified amino acids were analyzed using a ZipChip CE system (908 Devices, Boston, MA, USA) coupled to an Orbitrap Fusion Tribrid mass spectrometer (Thermo Fisher Scientific). The obtained data were analyzed using the Qual Browser in Xcalibur (Thermo Fisher Scientific) version 4.3.73.11. To identify carbon fixation pathways, we compared the observed mass fractions with predicted labeling structures from genomic information in KEGG PATHWAY Database^51^ and MetaCyc^52^.

## Acknowledgments

Part of this work was supported by the Green Innovation Fund Project, JPNP22010, commissioned by the New Energy and Industrial Technology Development Organization (NEDO) and Grant-in-Aid for Scientific Research (22H05152), (21K15024), (23H04656), and (23H04654) from the Ministry of Education, Culture, Sports, Science and Technology (MEXT). We thank Eiji Tasumi for his support in setting up the gas measurements. Bioinformatics analysis was partially performed using the supercomputing system at the National Institute of Genetics (NIG), Research Organization of Information and Systems (ROIS), and the Earth Simulator systems at JAMSTEC.

## Author contributions

Y.F., T.N., and K.T. designed the research study. S.Sanae, Y.C, and T.W, cultivated the strains. Y.F. and S.Shimamura performed metabolomics and proteomics. Y.F. and S.H performed bioinformatic analysis. Y.F. wrote the original draft. All authors reviewed the manuscript and approved the final manuscript.

## Data availability

The complete genome sequences of *Archaeoglobus* sp. strain MCR have been deposited in the DNA Data Bank of Japan under the accession number AP039546.1. All LC– MS/MS raw files of the proteome analysis and complete protein identification lists were deposited in the ProteomeXchange Consortium via jPOSTrepo^53^ (Okuda et al., 2025), under the data set identifiers PXD066747 (JPST003963) for *Archaeoglobus* sp. strain MCR. CE–MS data are included in the published article or its supplementary information files. Raw CE–MS data are available from the corresponding author upon request.

## Supplementary Information is available for this paper

