## Supplemental Information for "Direct carbon monoxide fixation via the bacterial and archaeal Wood–Ljungdahl pathways"

### Supplementary Methods

#### Growth characterization of *Archaeoglobus* sp. strain MCR under carboxydotrophic conditions

For growth characterization of *Archaeoglobus* sp. strain MCR (=DSM 106437), the archaeon was grown at 75 °C on 20 mL of basal medium. The basal medium, which was slightly modified MJ synthetic seawater[1] contained the following components per liter: 0.5 g $NH_4Cl$, 0.14 g $K_2HPO_4$, 0.33 g KCl, 4.18 g $MgCl_2{\cdot}6H_2O$, 3.4 g $MgSO_4{\cdot}6H_2O$, 0.14 g $CaCl_2{\cdot}2H_2O$, 20 g NaCl, 2g $NaSO_4$, 5 mg $NiCl_2{\cdot}6H_2O$, 2 mg $Fe(NH_4)_2(SO_4)_2{\cdot}6H_2O$, 1 mL trace mineral solution, 1 mL selenite-tungstate solution[2], 1 mL vitamin solution (ten times stock solution of Balch's vitamin solution[3], 1 mg resazurin, 1 g $NaHCO_3$, and 0.5 g $Na_2S{\cdot}9H_2O$. The trace mineral solution composed 3.8 g/L $FeCl_2$, 0.83g/L $CoCl_2$, 5.3 g/L $MnCl_2$ $4H_2O$, 0.85 g/L $ZnCl_2$, 0.1 $H_3BO_3$, 2.6 g/L $NiCl_2$, 0.01 g/L $AlCl_3$, 0.1 g/L $Na_2MoO_4$ $2H_2O$ and 0.05 g/L $CuCl_2$. The gas phase of the inoculated vessels was changed to a mixture of 20%$CO_2$, 50%CO, and 30%$N_2$ (0.3 MPa). Growth was assessed by observation under an epifluorescence microscope (BX53, Olympus). The CO, $H_2$, and $CO_2$ concentrations were monitored using a gas chromatograph 7890B GC system with a pulsed discharge helium ionization detector (Agilent Technologies).

#### Phylogenetic analysis of carbon monoxide dehydrogenases (CODHs) and their metabolic function prediction

Five CODHs of *Archaeoglobus* sp. strain MCR (GenBank record AP039546.1) were predicted using BLASTp[4] with default settings. For the similarity-based search, CODHs from *Carboxydothermus hydrogenoformans* (WP_011343033; CooS-type) and *Methanosarcina barkeri* (WP_011305243; Cdh-type) were used, as previously described[5,

[6, 7]. In addition, we used the comprehensive non-redundant CODH sequence database[7] to retrieve 2462 sequences, including two CODHs of *T. indicus* strain CIR29812$^T$ (=DSM 15286$^T$=JCM 11889$^T$). A total of 2467 CODH sequences were aligned using MAFFT[8] with '--auto' settings. A phylogenetic tree was reconstructed using FastTree[9] with the JTT + CAT model. The phylogenetic clades and metabolic functions were estimated based on phylogenetic placement and genomic context, as previously described[5, 6, 7].

**Protein extraction for proteomic analysis**

For proteomic analysis of *Archaeoglobus* sp. strain MCR, 20 mL culture at the exponential phase was harvested by centrifugation (4°C, 15,000 rpm, 5 min), and then the cell pellet was stored at −80°C until use. The stored cell pellets were suspended in 100 mM triethylammonium bicarbonate (TEAB) buffer containing 2 mM phenylmethylsulfonyl fluoride, followed by disruption by sonication. The dried proteins were obtained by evaporation and then denatured with MPEX PTS Reagent Solutions (GL Science, Tokyo, Japan). After alkylation using dithiothreitol and iodoacetamide, protein digestion was initiated by trypsin protease, followed by the removal of detergents in the samples. The digested protein solutions were evaporated to dryness, and then the dried proteins were resuspended in water containing 2% acetonitrile and 0.1% trifluoroacetic acid for proteomic analysis.

**Natural abundance of $^{13}C$ at $^{13}C$ tracer-based metabolomics**

In cells grown with non-labeled substrates, isotopologues of natural $^{13}C$ were detected in mass fractions of M+1. Generally, the relative abundance of naturally $^{13}C$-labeled carbon is proportional to the number of carbon atoms composed in each fragment derived from amino acids. Thus, the incorporation rate of $^{13}C$ in vivo labeling was evaluated after eliminating the natural abundance of $^{13}C$ (1.1%), as previously described[10].
