## Supplemental Figures for "Direct carbon monoxide fixation via the bacterial and archaeal Wood–Ljungdahl pathways"

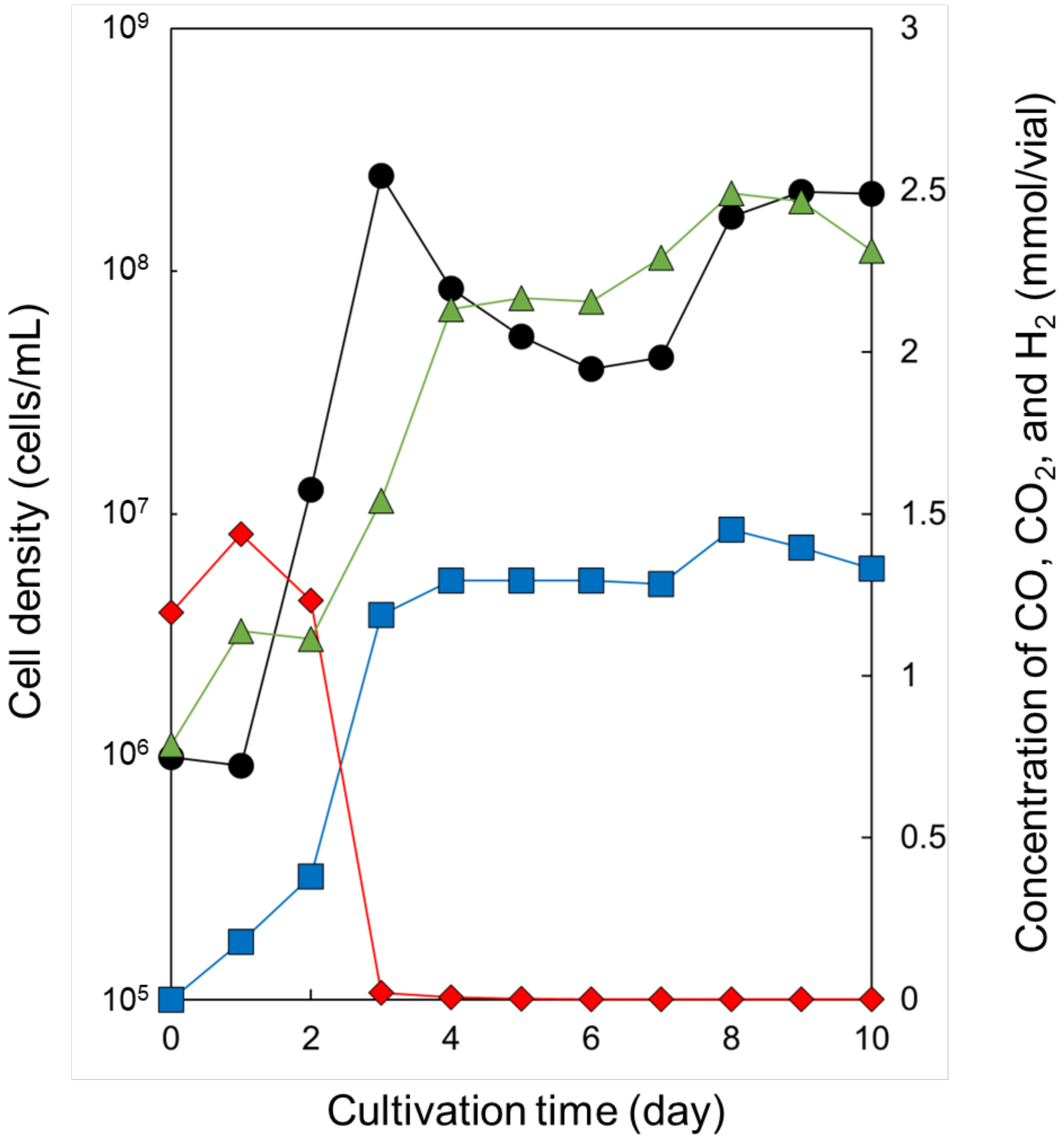


**Fig. S1 Growth of *Archaeoglobus* sp. strain MCR under the carboxydotrophic conditions**

Different parameters are shown: black disks, cell density; red diamonds, CO concentration; green triangles, $CO_2$ concentration; blue squares, $H_2$ concentration.

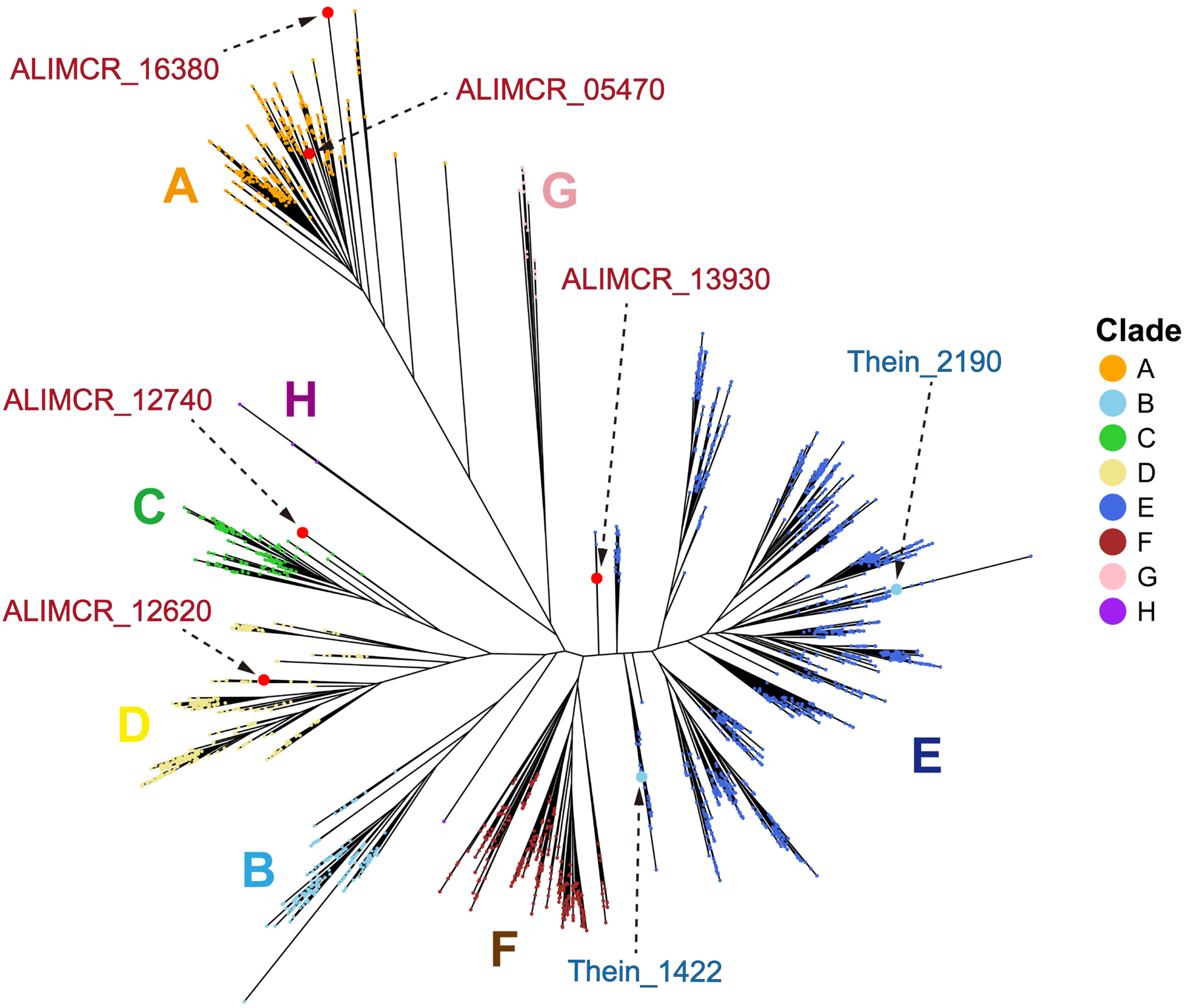


**Fig. S2 Phylogenetic tree of CODHs.**
Genes from *T. indicus* and *Archaeoglobus* sp. strain MCR are indicated in blue and red texts, respectively. Node color indicates phylogenetic clades defined previously (Inoue et al., 2022).

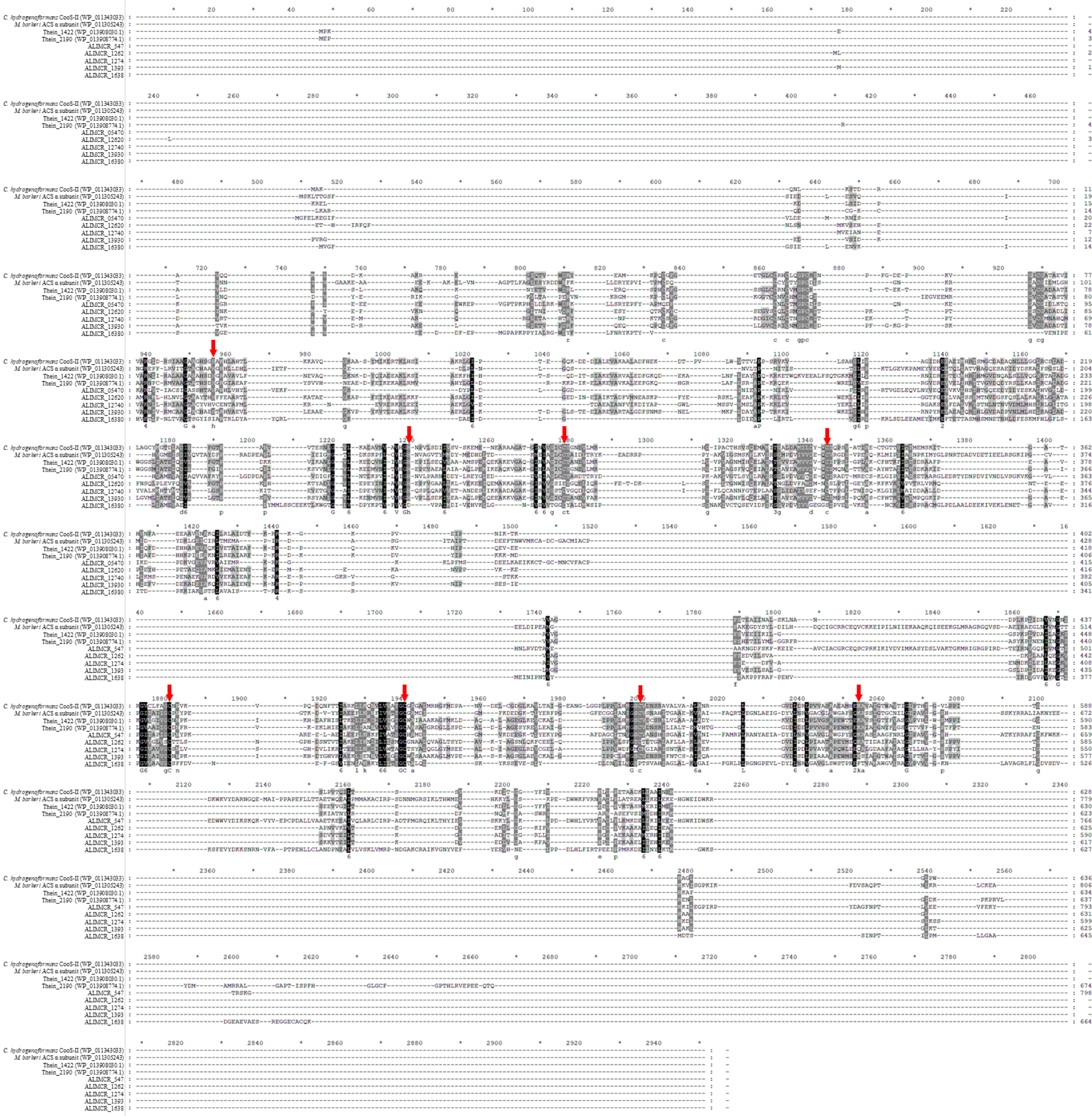

**Fig. S3 Multiple alignment of CODHs from *T. indicus* and *Archaeoglobus* sp. strain MCR**

Red arrows indicate the residues of active-site motifs in the Ni-Fe-S-type metallocluster (C-cluster) and acid-base around C-cluster

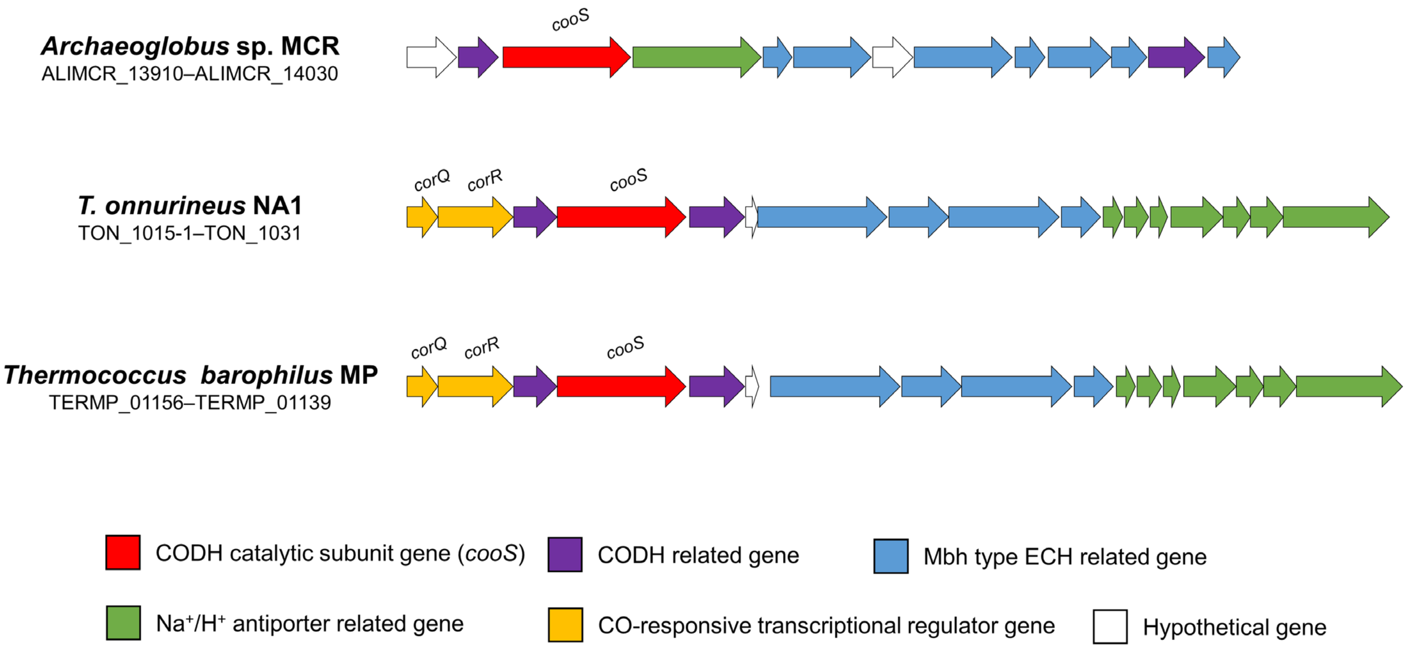


**Fig. S4 Comparison of the CODH−Mbh type ECH gene cluster among *Archaeoglobus* sp. strain MCR and strains in the genus *Thermococcus***

| Metabolism | gene ID | gene symbol | K number | Description | Relative abundance (%) | | | | Log$_2$ fold change CO/CO$_2$ + sulfate vs H$_2$/CO$_2$ + sulfate | Sum PEP Score |
|---|---|---|---|---|---|---|---|---|---|---|
| | | | | | CO/CO$_2$ | CO/CO$_2$ + sulfate | CO + sulfate | H$_2$/CO$_2$ + sulfate | | |
| **Carbon fixation** | | | | | | | | | | |
| **Imcomplete TCA cycle** | ALIMCR_12950 | *porC* | K00172 | pyruvate ferredoxin oxidoreductase gamma subunit | 69.7 | 100 | 70.9 | 36.4 | 1.46 | 36.4 |
| | ALIMCR 12960 | *porD* | K00171 | pyruvate ferredoxin oxidoreductase delta subunit | 43.9 | 54.0 | 100 | 21.7 | 1.32 | 17.0 |
| | ALIMCR 12970 | *porA* | K00169 | pyruvate ferredoxin oxidoreductase alpha subunit | 68.6 | 100 | 88.2 | 40.2 | 1.32 | 111.8 |
| | ALIMCR_12980 | *porB* | K00170 | pyruvate ferredoxin oxidoreductase beta subunit | 84.6 | 100 | 87.0 | 41.8 | 1.26 | 31.1 |
| | ALIMCR 20140 | *cs (gltA)* | K01647 | citrate synthase | 76.0 | 97.8 | 100 | 71.0 | 0.46 | 135.2 |
| | ALIMCR 04860 | *pycA* | K01959 | pyruvate carboxylase subunit A | 100 | 88.1 | 85.8 | 42.3 | 1.06 | 149.5 |
| | ALIMCR_06820 | *pycB* | K01960 | pyruvate carboxylase subunit B | 79.9 | 100 | 78.3 | 46.1 | 1.12 | 134.8 |
| | ALIMCR 483 | *IDH* | K00031 | isocitrate dehydrogenase | 86.6 | 94.0 | 100 | 70.3 | 0.42 | 224.5 |
| | ALIMCR 07440 | *sdhD* | K00242 | succinate dehydrogenase membrane anchor subunit | Not detected | | | | - | Not detected |
| | ALIMCR 07450 | *sdhC* | K00241 | succinate dehydrogenase cytochrome b subunit | Not detected | | | | - | Not detected |
| | ALIMCR_07460 | *sdhB* | K00240 | succinate dehydrogenase iron-sulfur subunit | Not detected | | | | - | Not detected |
| | ALIMCR 07470 | *sdhA* | K00239 | succinate dehydrogenase flavoprotein subunit | 46.5 | 88.3 | 73.0 | 100 | -0.18 | 48.0 |
| | ALIMCR 09260 | *fumB* | K01678 | fumarate hydratase subunit beta | 49.8 | 97.0 | 64.9 | 100 | -0.04 | 11.6 |
| | ALIMCR_09270 | *fumA* | K01677 | fumarate hydratase subunit alpha | 53.8 | 100 | 65.3 | 92.9 | 0.11 | 42.4 |
| | ALIMCR 06530 | *mdh* | K00024 | malate dehydrogenase | 68.8 | 100 | 69.0 | 36.2 | 1.47 | 103.8 |
| **WL pathway** | ALIMCR 07840 | *fwdC* | K00202 | formylmethanofuran dehydrogenase subunit C | 100 | 68.7 | 35.2 | 31.2 | 1.14 | 110.6 |
| | ALIMCR 07850 | *fwdA* | K00200 | formylmethanofuran dehydrogenase subunit A | 100 | 76.9 | 43.2 | 48.4 | 0.67 | 197.7 |
| | ALIMCR_07860 | *fwdB* | K00201 | formylmethanofuran dehydrogenase subunit B | 83.7 | 62.5 | 49.8 | 100 | -0.68 | 148.8 |
| | ALIMCR 07870 | *fwdD* | K00203 | formylmethanofuran dehydrogenase subunit D | 100 | 65.3 | 98.8 | 94.6 | -0.54 | 61.5 |
| | ALIMCR 19390 | *fwdF* | K00205 | 4Fe-4S ferredoxin | 62.2 | 61.6 | 100 | 61.6 | 0.00 | 32.4 |
| | ALIMCR_13250 | *ftr* | K00672 | formylmethanofuran--tetrahydromethanopterin N-formyltransferase | 81.7 | 88.2 | 100 | 84.4 | 0.06 | 103.9 |
| | ALIMCR 07020 | *mch* | K01499 | methenyltetrahydromethanopterin cyclohydrolase | 61.1 | 100 | 78.5 | 68.8 | 0.54 | 92.9 |
| | ALIMCR 05930 | *mtd* | K00319 | methylenetetrahydromethanopterin dehydrogenase | 88.6 | 48.2 | 100 | 59.9 | -0.31 | 395.0 |
| | ALIMCR 01800 | *mer* | K00320 | 5,10-methylenetetrahydromethanopterin reductase | 74.0 | 60.2 | 100 | 65.2 | -0.12 | 215.9 |
| **CODH–ACS gene cluster** | ALIMCR_05430 | - | K06940 | uncharacterized protein | Not detected | | | | - | Not detected |
| | ALIMCR_05440 | *cooF* | K11260 | 4Fe-4S ferredoxin | 92.3 | 100 | 80.2 | 79.6 | 0.33 | 9.5 |
| | ALIMCR_05450 | *fwdB* | K00201 | formylmethanofuran dehydrogenase subunit B | 100 | 86.9 | 88.5 | 70.4 | 0.31 | 87.6 |
| | ALIMCR_05460 | *fwdD* | K00203 | formylmethanofuran dehydrogenase subunit D | 70.5 | 90.0 | 100 | 82.6 | 0.12 | 24.4 |
| | ALIMCR_05470 | *cdhA* | K00192 | CODH/ACS complex subunit alpha | 60.8 | 44.4 | 97.9 | 100 | -1.17 | 600.2 |
| | ALIMCR_05480 | *cdhB* | K00195 | CODH/ACS complex subunit epsilon (Archaea specific) | 53.4 | 43.6 | 100 | 94.9 | -1.12 | 83.8 |
| | ALIMCR_05490 | *cdhC* | K00193 | CODH/ACS complex subunit beta | 61.7 | 54.8 | 76.8 | 100 | -0.87 | 513.9 |
| | ALIMCR_05500 | *cooC* | K07321 | CO dehydrogenase maturation factor | 49.3 | 62.9 | 85.4 | 100 | -0.67 | 88.4 |
| | ALIMCR_05510 | *cdhD* | K00194 | CODH/ACS complex subunit delta | 57.1 | 58.2 | 77.2 | 100 | -0.78 | 270.6 |
| | ALIMCR_05520 | *cdhE* | K00197 | CODH/ACS complex subunit gamma | 56.3 | 59.7 | 75.6 | 100 | -0.74 | 224.1 |
| **Energy metabolism** | | | | | | | | | | |
| **CODH–Mbh type gene cluster** | ALIMCR 13910 | - | K07013 | uncharacterized protein | Not detected | | | | - | Not detected |
| | ALIMCR_13920 | *cooF* | K00196 | anaerobic carbon-monoxide dehydrogenase iron sulfur subunit | 81.9 | 71.5 | 100 | 0.1 | 10.14 | 154.2 |
| | ALIMCR_13930 | *cooS* | K00198 | anaerobic carbon-monoxide dehydrogenase catalytic subunit | 98.6 | 83.3 | 100 | 0.2 | 8.62 | 949.3 |
| | ALIMCR_13940 | *mrpA* | K05565 | multicomponent Na$^+$:H$^+$ antiporter subunit A | 86.6 | 87.4 | 100 | 0.0 | >10 | 100.0 |
| | ALIMCR_13950 | *nuoCD* | K13378 | NADH-quinone oxidoreductase subunit C/D | 100 | 91.7 | 64.1 | 0.0 | >10 | 91.2 |
| | ALIMCR_13960 | *mbhL* | K18016 | membrane-bound hydrogenase subunit alpha | 100 | 94.2 | 65.3 | 0.5 | 7.69 | 217.5 |
| | ALIMCR_13970 | - | - | - | 100 | 60.7 | 14.3 | 0.0 | >10 | 18.9 |
| | ALIMCR_13980 | *fpoN* | K22169 | $F_{420}H_2$ dehydrogenase subunit N | 100 | 82.1 | 72.2 | 0.3 | 8.01 | 46.8 |
| | ALIMCR_13990 | *mbhJ* | K18023 | membrane-bound hydrogenase subunit mbhJ | 100 | 88.9 | 84.6 | 0.0 | >10 | 29.6 |
| | ALIMCR_14000 | *nuoH* | K00337 | NADH-quinone oxidoreductase subunit H | 90.1 | 70.1 | 100 | 0.0 | >10 | 37.8 |
| | ALIMCR_14010 | *hyfH* | K12143 | hydrogenase-4 component H | 100 | 98.9 | 80.6 | 1.5 | 6.00 | 97.3 |
| | ALIMCR_14020 | *cooC* | K07321 | CO dehydrogenase maturation factor | 61.4 | 100 | 88.0 | 0.6 | 7.32 | 140.7 |
| | ALIMCR_14030 | *hycI* | K08315 | hydrogenase 3 maturation protease | Not detected | | | | - | Not detected |
| **Hydrogenase** | ALIMCR_10790 | *hdrA* | K03388 | heterodisulfide reductase subunit A2 | 16.4 | 40.7 | 59.8 | 100 | -1.30 | 354.7 |
| | ALIMCR 10800 | *hdrC* | K03390 | heterodisulfide reductase subunit C2 | 14.8 | 41.1 | 56.6 | 100 | -1.28 | 77.6 |
| | ALIMCR 10810 | *hdrB* | K03389 | heterodisulfide reductase subunit B2 | 17.2 | 36.9 | 44.4 | 100 | -1.44 | 122.6 |
| | ALIMCR_10820 | *vhuD* | K14127 | $F_{420}$-non-reducing hydrogenase iron-sulfur subunit | 14.6 | 51.0 | 67.5 | 100 | -0.97 | 66.0 |
| | ALIMCR_10830 | *vhuG* | K14128 | $F_{420}$-non-reducing hydrogenase small subunit | 16.5 | 33.1 | 43.1 | 100 | -1.59 | 142.4 |
| | ALIMCR_10840 | *vhuA* | K14126 | $F_{420}$-non-reducing hydrogenase large subunit | 16.9 | 35.9 | 44.3 | 100 | -1.48 | 369.0 |
| | ALIMCR_10850 | *vhtD-1* | K03605 | hydrogenase maturation protease | 100 | 98.1 | 25.0 | 14.6 | 2.75 | 11.1 |
| | ALIMCR 16470 | *vhtG* | K06282 | methanophenazine hydrogenase | 33.9 | 66.7 | 81.3 | 100 | -0.58 | 85.0 |
| | ALIMCR 16480 | *vhtA* | - | methanophenazine hydrogenase, large subunit | 36.2 | 81.9 | 70.8 | 100 | -0.29 | 266.3 |
| | ALIMCR_16490 | *vhtC* | K14069 | methanophenazine hydrogenase, cytochrome b subunit | 6.8 | 0.0 | 100 | 0.0 | >10 | 7.2 |
| | ALIMCR 16500 | *vhtD-2* | - | - | Not detected | | | | - | Not detected |
| **Dissimilatory sulfur redction metabolism** | ALIMCR 08710 | *dsrD* | - | - | 54.3 | 100 | 58.8 | 79.9 | 0.32 | 57.7 |
| | ALIMCR_08720 | *dsrB* | K11181 | dissimilatory sulfite reductase beta subunit | 43.1 | 68.8 | 57.8 | 100 | -0.54 | 298.7 |
| | ALIMCR_08730 | *dsrA* | K11180 | dissimilatory sulfite reductase alpha subunit | 49.5 | 73.2 | 61.2 | 100 | -0.45 | 241.1 |
| | ALIMCR_10690 | *dsrJ* | K27189 | [DsrC]-trisulfide reductase subunit J | 0.0 | 43.2 | 50.1 | 100 | -1.21 | 2.5 |
| | ALIMCR_10700 | *dsrK* | K27188 | [DsrC]-trisulfide reductase subunit K | 3.3 | 51.6 | 61.2 | 100 | -0.95 | 297.0 |
| | ALIMCR_10710 | *dsrM* | K27187 | [DsrC]-trisulfide reductase subunit M | 1.4 | 37.0 | 59.3 | 100 | -1.43 | 70.9 |
| | ALIMCR_10720 | *dsrP* | K27191 | [DsrC]-trisulfide reductase subunit P | 0.7 | 26.0 | 32.8 | 100 | -1.94 | 28.2 |
| | ALIMCR_10730 | *dsrO* | K27190 | [DsrC]-trisulfide reductase subunit O | 9.6 | 42.0 | 50.4 | 100 | -1.25 | 174.7 |
| | ALIMCR_10740 | *dsvC(dsrC)* | K23077 | dissimilatory sulfite reductase related protein | 62.2 | 71.5 | 81.8 | 100 | -0.48 | 222.1 |
| | ALIMCR_11370 | *sat* | K00958 | sulfate adenylyltransferase | 48.6 | 44.7 | 51.5 | 100 | -1.16 | 633.5 |
| | ALIMCR_11390 | *aprB* | K00395 | adenylylsulfate reductase, subunit B | 59.9 | 55.6 | 87.4 | 100 | -0.85 | 203.2 |
| | ALIMCR_11400 | *aprA* | K00394 | adenylylsulfate reductase, subunit A | 44.1 | 50.3 | 52.9 | 100 | -0.99 | 674.4 |
| | ALIMCR_20770 | ppx1 | K15986 | manganese-dependent inorganic pyrophosphatase | 71.2 | 49.6 | 53.5 | 100 | -1.01 | 369.6 |

Relative abundance (%): 0 – 100

Sum PEP score: 0 – 949.3

**Fig. S5 Changes of protein levels involved in genes for carbon fixation and energy metabolism from proteomic analysis in *Archaeoglobus sp.* strain MCR grown under chemolithoautotrophic or carboxydotrophic conditions**

Relative values of protein expression among 4 tested conditions are shown as percentages indicating abundance, where “100%” indicates the highest protein expression among all conditions. The orange color becomes deeper as the relative abundance values increase. Sum PEP scores indicate the relative values of protein amounts for the comparison of protein expression levels between different CDSs. The green color becomes deeper as the Sum PEP scores increase.

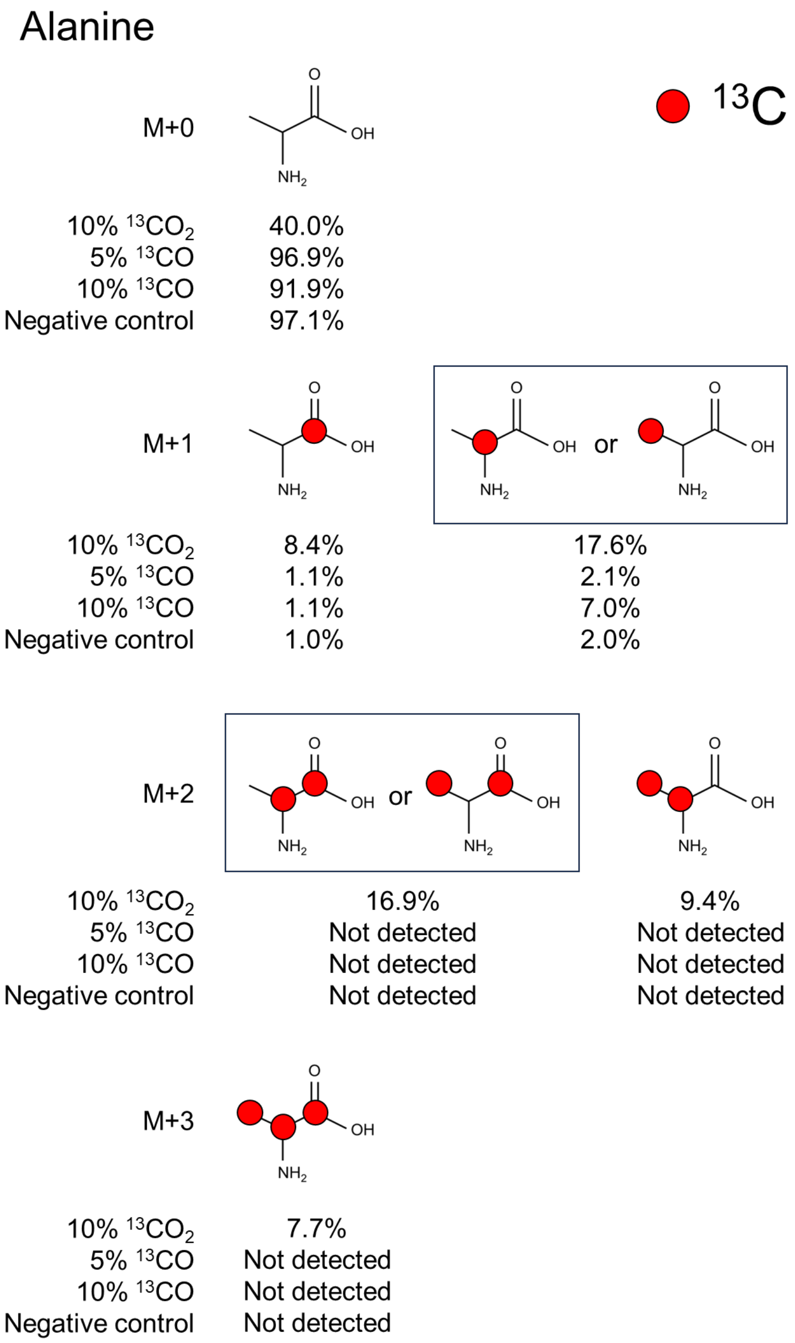


| | Isotopomer pattern | | Natural abundances of isotopes | *T. indicus* | | | |
|---|---|---|---|---|---|---|---|
| | No. of $^{13}C$ | | | 10% $^{13}CO_2$ | 5% $^{13}CO$ | 10% $^{13}CO$ | Negative control |
| Amino acid | C-1 | Others | Abundance (%) | Abundance (%) | Abundance (%) | Abundance (%) | Abundance (%) |
| $^{13}C_0$ Ala | 0 | 0 | 96.7 | 40.0 | 96.9 | 91.9 | 97.1 |
| $^{13}C_1$ Ala | 1 | 0 | 1.1 | 8.4 | 1.1 | 1.1 | 1.0 |
| | 0 | 1 | 2.2 | 17.6 | 2.1 | 7.0 | 2.0 |
| $^{13}C_2$ Ala | 1 | 1 | 0.0 | 16.9 | N.D. | N.D. | N.D. |
| | 0 | 2 | 0.0 | 9.4 | N.D. | N.D. | N.D. |
| $^{13}C_3$ Ala | 1 | 2 | 0.0 | 7.7 | N.D. | N.D. | N.D. |

N.D., not detected

**Fig. S6 The relative abundance of isotopomer pattern in Ala from *T. indicus***

The mass fractions for M+0, M+1, M+2, and M+3 represent fragments containing 0 to 3 $^{13}C$-labeled carbons, respectively. Red circles show $^{13}C$-labeled carbon atoms in the carbon skeleton.

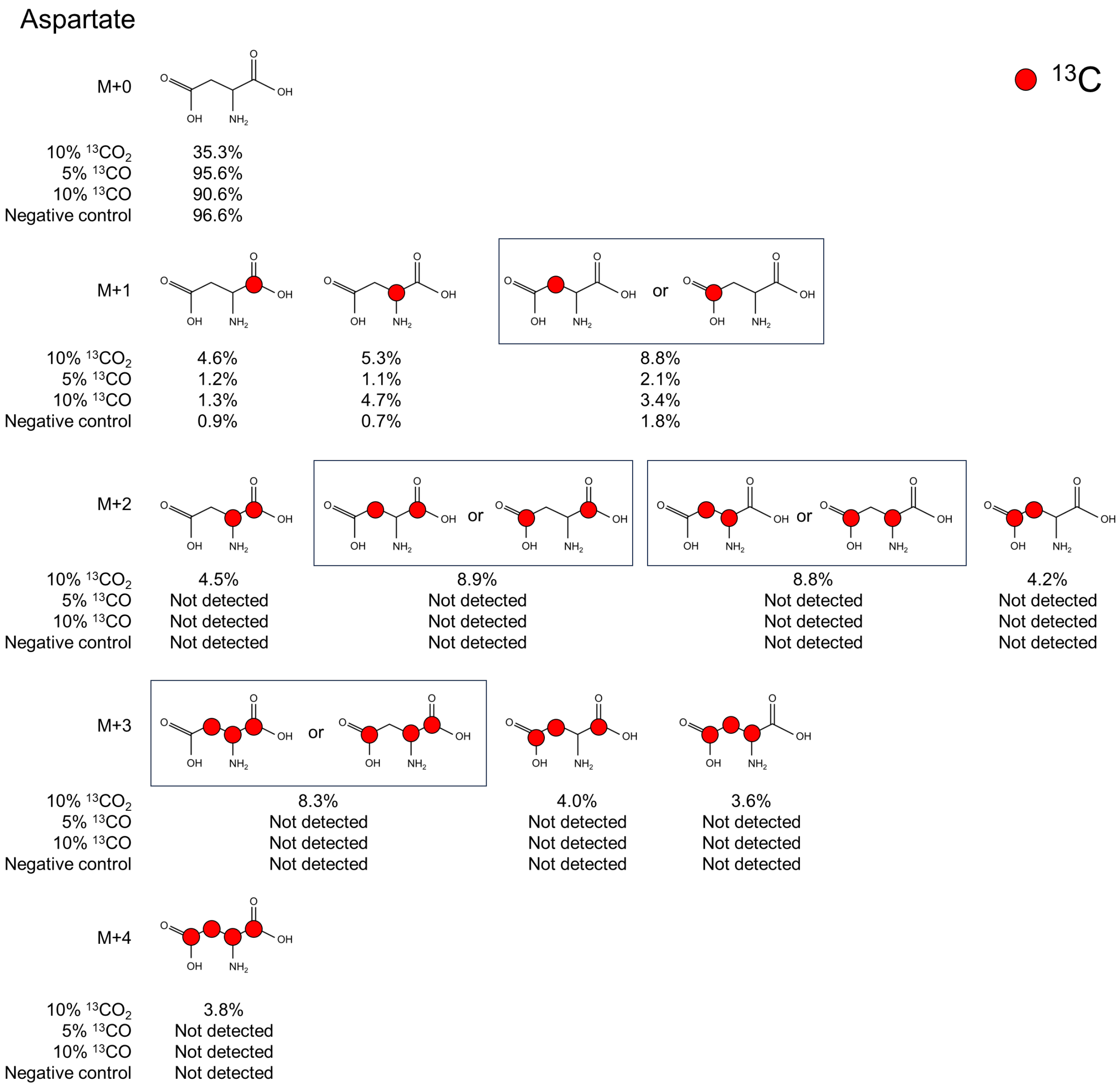


| | Isotopomer pattern | | Natural abundances of isotopes | *T. indicus* | | | |
|---|---|---|---|---|---|---|---|
| | No. of $^{13}C$ | | | 10% $^{13}CO_2$ | 5% $^{13}CO$ | 10% $^{13}CO$ | Negative control |
| **Amino acid** | **C-1** | **Others** | **abundance (%)** | **Abundance (%)** | **Abundance (%)** | **Abundance (%)** | **Abundance (%)** |
| $^{13}C_0$ Asp | 0 | 0 | 95.7 | 35.3 | 95.6 | 90.6 | 96.6 |
| $^{13}C_1$ Asp | 1 | 0 | 1.1 | 4.6 | 1.2 | 1.3 | 0.9 |
| | 0 | 1 | 3.2 | 14.1 | 3.2 | 8.1 | 2.5 |
| $^{13}C_2$ Asp | 1 | 1 | 0.0 | 13.4 | N.D. | N.D. | N.D. |
| | 0 | 2 | 0.0 | 13.0 | N.D. | N.D. | N.D. |
| $^{13}C_3$ Asp | 1 | 2 | 0.0 | 12.3 | N.D. | N.D. | N.D. |
| | 0 | 3 | 0.0 | 3.6 | N.D. | N.D. | N.D. |
| $^{13}C_4$ Asp | 1 | 3 | 0.0 | 3.8 | N.D. | N.D. | N.D. |
| **Amino acid** | **C-1, 2** | **C-3, 4** | **Abundance (%)** | **Abundance (%)** | **Abundance (%)** | **Abundance (%)** | **Abundance (%)** |
| $^{13}C_0$ Asp | 0 | 0 | 95.7 | 35.3 | 95.6 | 90.6 | 96.6 |
| $^{13}C_1$ Asp | 1 | 0 | 2.1 | 9.9 | 2.3 | 6.0 | 1.6 |
| | 0 | 1 | 2.1 | 8.8 | 2.1 | 3.4 | 1.8 |
| $^{13}C_2$ Asp | 2 | 0 | 0.0 | 4.5 | N.D. | N.D. | N.D. |
| | 1 | 1 | 0.0 | 17.7 | N.D. | N.D. | N.D. |
| | 0 | 2 | 0.0 | 4.2 | N.D. | N.D. | N.D. |
| $^{13}C_3$ Asp | 2 | 1 | 0.0 | 8.3 | N.D. | N.D. | N.D. |
| | 1 | 2 | 0.0 | 7.6 | N.D. | N.D. | N.D. |
| $^{13}C_4$ Asp | 2 | 2 | 0.0 | 3.8 | N.D. | N.D. | N.D. |

N.D., not detected

**Fig. S7 The relative abundance of isotopomer pattern in Asp from *T. indicus***

The mass fractions for M+0, M+1, M+2, M+3, and M+4 represent fragments containing 0 to 4 $^{13}C$-labeled carbons, respectively. Red circles show $^{13}C$-labeled carbon atoms in the carbon skeleton.

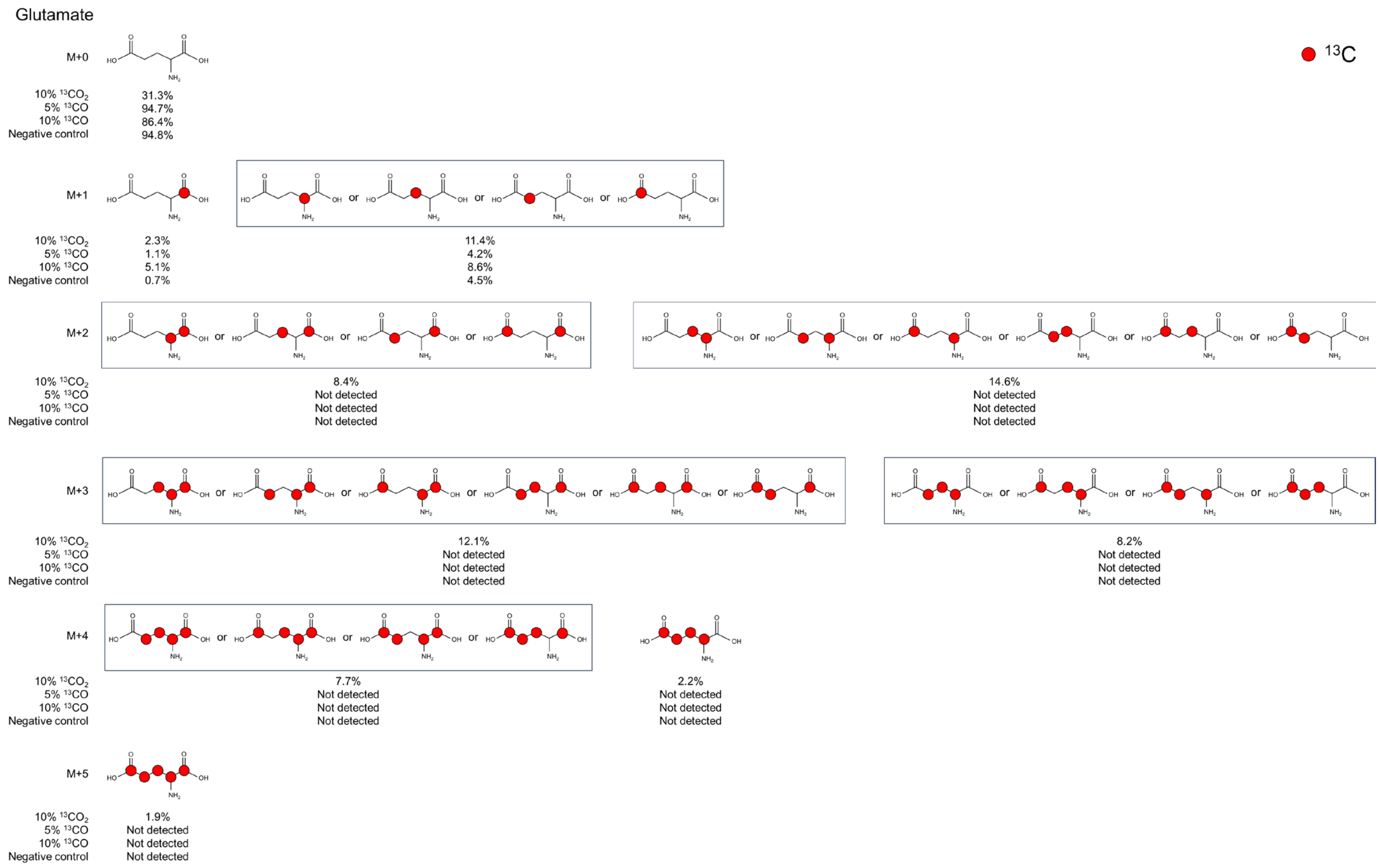


| | Isotopomer pattern | | Natural abundances of isotopes | *T. indicus* | | | |
|---|---|---|---|---|---|---|---|
| | No. of $^{13}$C | | | 10% $^{13}CO_2$ | 5% $^{13}$CO | 10% $^{13}$CO | Negative control |
| Amino acid | C-1 | Others | Abundance (%) | Abundance (%) | Abundance (%) | Abundance (%) | Abundance (%) |
| $^{13}C_0$ Glu | 0 | 0 | 94.6 | 31.3 | 94.7 | 86.4 | 94.8 |
| $^{13}C_1$ Glu | 1 | 0 | 1.1 | 2.3 | 1.1 | 5.1 | 0.7 |
| | 0 | 1 | 4.2 | 11.4 | 4.2 | 8.6 | 4.5 |
| $^{13}C_2$ Glu | 1 | 1 | 0.0 | 8.4 | N.D. | N.D. | N.D. |
| | 0 | 2 | 0.1 | 14.6 | N.D. | N.D. | N.D. |
| $^{13}C_3$ Glu | 1 | 2 | 0.0 | 12.1 | N.D. | N.D. | N.D. |
| | 0 | 3 | 0.0 | 8.2 | N.D. | N.D. | N.D. |
| $^{13}C_4$ Glu | 1 | 3 | 0.0 | 7.7 | N.D. | N.D. | N.D. |
| | 0 | 4 | 0.0 | 2.2 | N.D. | N.D. | N.D. |
| $^{13}C_5$ Glu | 1 | 4 | 0.0 | 1.9 | N.D. | N.D. | N.D. |

N.D., not detected

**Fig. S8 The relative abundance of isotopomer pattern in Glu from *T. indicus***

The mass fractions for M+0, M+1, M+2, M+3, M+4, and M+5 represent fragments containing 0 to 5 $^{13}$C-labeled carbons, respectively. Red circles show $^{13}$C-labeled carbon atoms in the carbon skeleton.

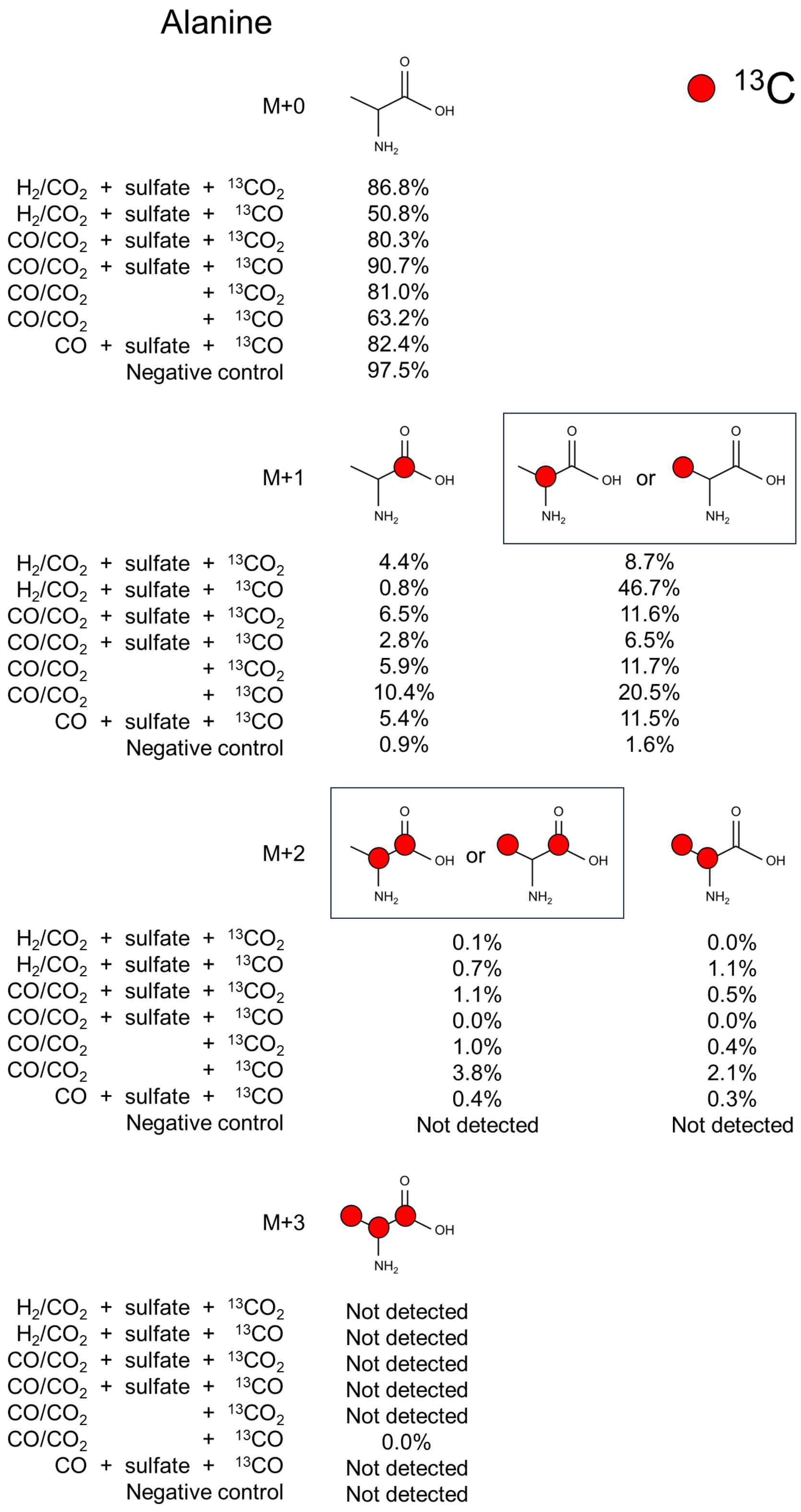


| | Isotopomer pattern | | Natural abundances of isotopes | *A. lithotrophicus* strain MCR | | | | | | | |
|---|---|---|---|---|---|---|---|---|---|---|---|
| | | | | $H_2/CO_2$ | | CO/$CO_2$ | | | | CO | Negative control |
| | No. of $^{13}C$ | | | $^{13}CO_2$ + sulfate | $^{13}CO$ + sulfate | $^{13}CO_2$ + sulfate | $^{13}CO$ + sulfate | $^{13}CO_2$ | $^{13}CO$ | $^{13}CO$ + sulfate | |
| Amino acid | C-1 | Others | Abundance (%) | Abundance (%) | Abundance (%) | Abundance (%) | Abundance (%) | Abundance (%) | Abundance (%) | Abundance (%) | Abundance (%) |
| $^{13}C_0$ Ala | 0 | 0 | 96.7 | 86.8 | 50.8 | 80.3 | 90.7 | 81.0 | 63.2 | 82.4 | 97.5 |
| $^{13}C_1$ Ala | 1 | 0 | 1.1 | 4.4 | 0.8 | 6.5 | 2.8 | 5.9 | 10.4 | 5.4 | 0.9 |
| | 0 | 1 | 2.2 | 8.7 | 46.7 | 11.6 | 6.5 | 11.7 | 20.5 | 11.5 | 1.6 |
| $^{13}C_2$ Ala | 1 | 1 | 0.0 | 0.1 | 0.7 | 1.1 | 0.0 | 1.0 | 3.8 | 0.4 | N.D. |
| | 0 | 2 | 0.0 | 0.0 | 1.1 | 0.5 | 0.0 | 0.4 | 2.1 | 0.3 | N.D. |
| $^{13}C_3$ Ala | 1 | 2 | 0.0 | N.D. | N.D. | N.D. | N.D. | N.D. | 0.0 | N.D. | N.D. |

N.D., not detected

**Fig. S9 The relative abundance of isotopomer pattern in Ala from *Archaeoglobus* sp. strain MCR**

The mass fractions for M+0, M+1, M+2, and M+3 represent fragments containing 0 to 3 $^{13}C$-labeled carbons, respectively. Red circles show $^{13}C$-labeled carbon atoms in the carbon skeleton.

Aspartate

● $^{13}C$

M+0

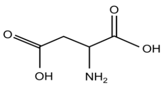

| | |
|---|---|
| $H_2/CO_2$ + sulfate + $^{13}CO_2$ | 80.9% |
| $H_2/CO_2$ + sulfate + $^{13}CO$ | 49.4% |
| $CO/CO_2$ + sulfate + $^{13}CO_2$ | 73.2% |
| $CO/CO_2$ + sulfate + $^{13}CO$ | 87.4% |
| $CO/CO_2$ + $^{13}CO_2$ | 75.0% |
| $CO/CO_2$ + $^{13}CO$ | 54.4% |
| CO + sulfate + $^{13}CO$ | 76.6% |
| Negative control | 96.2% |

M+1

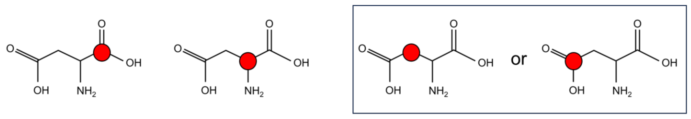

| | | | |
|---|---|---|---|
| $H_2/CO_2$ + sulfate + $^{13}CO_2$ | 4.1% | 3.8% | 9.2% |
| $H_2/CO_2$ + sulfate + $^{13}CO$ | 0.7% | 41.2% | 5.5% |
| $CO/CO_2$ + sulfate + $^{13}CO_2$ | 6.4% | 5.3% | 11.6% |
| $CO/CO_2$ + sulfate + $^{13}CO$ | 2.6% | 2.8% | 6.4% |
| $CO/CO_2$ + $^{13}CO_2$ | 6.6% | 5.0% | 10.4% |
| $CO/CO_2$ + $^{13}CO$ | 9.5% | 7.6% | 17.5% |
| CO + sulfate + $^{13}CO$ | 5.3% | 4.7% | 10.7% |
| Negative control | 0.9% | 1.2% | 1.8% |

M+2

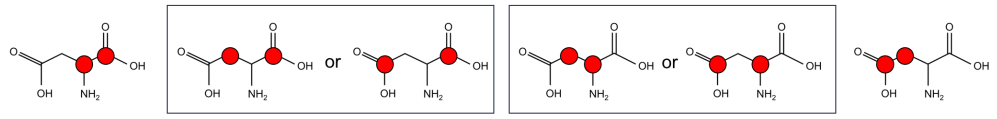

| | | | | |
|---|---|---|---|---|
| $H_2/CO_2$ + sulfate + $^{13}CO_2$ | 0.2% | 0.8% | 0.1% | 0.9% |
| $H_2/CO_2$ + sulfate + $^{13}CO$ | 0.6% | 0.3% | 1.9% | 0.3% |
| $CO/CO_2$ + sulfate + $^{13}CO_2$ | 0.3% | 1.2% | 0.9% | 1.2% |
| $CO/CO_2$ + sulfate + $^{13}CO$ | 0.4% | 0.3% | Not detected | 0.1% |
| $CO/CO_2$ + $^{13}CO_2$ | 0.3% | 1.4% | 0.5% | 1.0% |
| $CO/CO_2$ + $^{13}CO$ | 1.0% | 3.7% | 2.0% | 2.9% |
| CO + sulfate + $^{13}CO$ | 0.2% | 1.2% | 0.3% | 1.0% |
| Negative control | Not detected | Not detected | Not detected | Not detected |

M+3

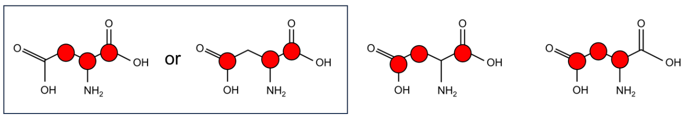

| | | | |
|---|---|---|---|
| $H_2/CO_2$ + sulfate + $^{13}CO_2$ | Not detected | Not detected | Not detected |
| $H_2/CO_2$ + sulfate + $^{13}CO$ | Not detected | Not detected | Not detected |
| $CO/CO_2$ + sulfate + $^{13}CO_2$ | Not detected | Not detected | Not detected |
| $CO/CO_2$ + sulfate + $^{13}CO$ | Not detected | Not detected | Not detected |
| $CO/CO_2$ + $^{13}CO_2$ | Not detected | Not detected | Not detected |
| $CO/CO_2$ + $^{13}CO$ | 0.6% | 0.6% | Not detected |
| CO + sulfate + $^{13}CO$ | Not detected | 0.1% | Not detected |
| Negative control | Not detected | Not detected | Not detected |

M+4

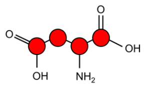

| | |
|---|---|
| $H_2/CO_2$ + sulfate + $^{13}CO_2$ | Not detected |
| $H_2/CO_2$ + sulfate + $^{13}CO$ | Not detected |
| $CO/CO_2$ + sulfate + $^{13}CO_2$ | Not detected |
| $CO/CO_2$ + sulfate + $^{13}CO$ | Not detected |
| $CO/CO_2$ + $^{13}CO_2$ | Not detected |
| $CO/CO_2$ + $^{13}CO$ | 0.1% |
| CO + sulfate + $^{13}CO$ | Not detected |
| Negative control | Not detected |

| | Isotopomer pattern | | Natural abundances of isotopes | *A. lithotrophicus* strain MCR | | | | | | | |
|---|---|---|---|---|---|---|---|---|---|---|---|
| | No. of $^{13}C$ | | | $H_2/CO_2$ | | $CO/CO_2$ | | | | CO | Negative control |
| | | | | $^{13}CO_2$ + sulfate | $^{13}CO$ + sulfate | $^{13}CO_2$ + sulfate | $^{13}CO$ + sulfate | $^{13}CO_2$ | $^{13}CO$ | $^{13}CO$ + sulfate | |
| **Amino acid** | **C-1** | **Others** | **Abundance (%)** | **Abundance (%)** | **Abundance (%)** | **Abundance (%)** | **Abundance (%)** | **Abundance (%)** | **Abundance (%)** | **Abundance (%)** | **Abundance (%)** |
| $^{13}C_0$ Asp | 0 | 0 | 95.7 | 80.9 | 49.4 | 73.2 | 87.4 | 75.0 | 54.4 | 76.6 | 96.2 |
| $^{13}C_1$ Asp | 1 | 0 | 1.1 | 4.1 | 0.7 | 6.4 | 2.6 | 6.6 | 9.5 | 5.3 | 0.9 |
| | 0 | 1 | 3.2 | 13.0 | 46.7 | 16.9 | 9.2 | 15.3 | 25.0 | 15.4 | 2.9 |
| $^{13}C_2$ Asp | 1 | 1 | 0.0 | 1.0 | 0.9 | 1.5 | 0.7 | 1.7 | 4.7 | 1.4 | N.D. |
| | 0 | 2 | 0.0 | 1.0 | 2.2 | 2.0 | 0.0 | 1.4 | 4.9 | 1.3 | N.D. |
| $^{13}C_3$ Asp | 1 | 2 | 0.0 | N.D. | N.D. | N.D. | N.D. | N.D. | 1.3 | N.D. | N.D. |
| | 0 | 3 | 0.0 | N.D. | N.D. | N.D. | N.D. | N.D. | N.D. | N.D. | N.D. |
| $^{13}C_4$ Asp | 1 | 3 | 0.0 | N.D. | N.D. | N.D. | N.D. | N.D. | 0.1 | N.D. | N.D. |
| **Amino acid** | **C-1, 2** | **C-3, 4** | **Abundance (%)** | **Abundance (%)** | **Abundance (%)** | **Abundance (%)** | **Abundance (%)** | **Abundance (%)** | **Abundance (%)** | **Abundance (%)** | **Abundance (%)** |
| $^{13}C_0$ Asp | 0 | 0 | 95.7 | 80.9 | 49.4 | 73.2 | 87.4 | 75.0 | 54.4 | 76.6 | 96.2 |
| $^{13}C_1$ Asp | 1 | 0 | 2.1 | 8.0 | 42.0 | 11.7 | 5.4 | 11.5 | 17.1 | 10.0 | 2.1 |
| | 0 | 1 | 2.1 | 9.2 | 5.5 | 11.6 | 6.4 | 10.4 | 17.5 | 10.7 | 1.8 |
| $^{13}C_2$ Asp | 2 | 0 | 0.0 | 0.2 | 0.6 | 0.3 | 0.4 | 0.3 | 1.0 | 0.2 | N.D. |
| | 1 | 1 | 0.0 | 0.9 | 2.2 | 2.1 | 0.2 | 1.9 | 5.7 | 1.4 | N. D. |
| | 0 | 2 | 0.0 | 0.9 | 0.3 | 1.2 | 0.1 | 1.0 | 2.9 | 1.0 | N.D. |
| $^{13}C_3$ Asp | 2 | 1 | 0.0 | N.D. | N.D. | N.D. | N.D. | N.D. | 0.6 | N.D. | N.D. |
| | 1 | 2 | 0.0 | N.D. | N.D. | N.D. | N.D. | N.D. | 0.6 | 0.1 | N.D. |
| $^{13}C_4$ Asp | 2 | 2 | 0.0 | N.D. | N.D. | N.D. | N.D. | N.D. | 0.1 | N.D. | N.D. |

N.D., not detected

**Fig. S10 The relative abundance of isotopomer pattern in Asp from *Archaeoglobus* sp. strain MCR**

The mass fractions for M+0, M+1, M+2, M+3, and M+4 represent fragments containing 0 to 4 $^{13}C$-labeled carbons, respectively. Red circles show $^{13}C$-labeled carbon atoms in the carbon skeleton.

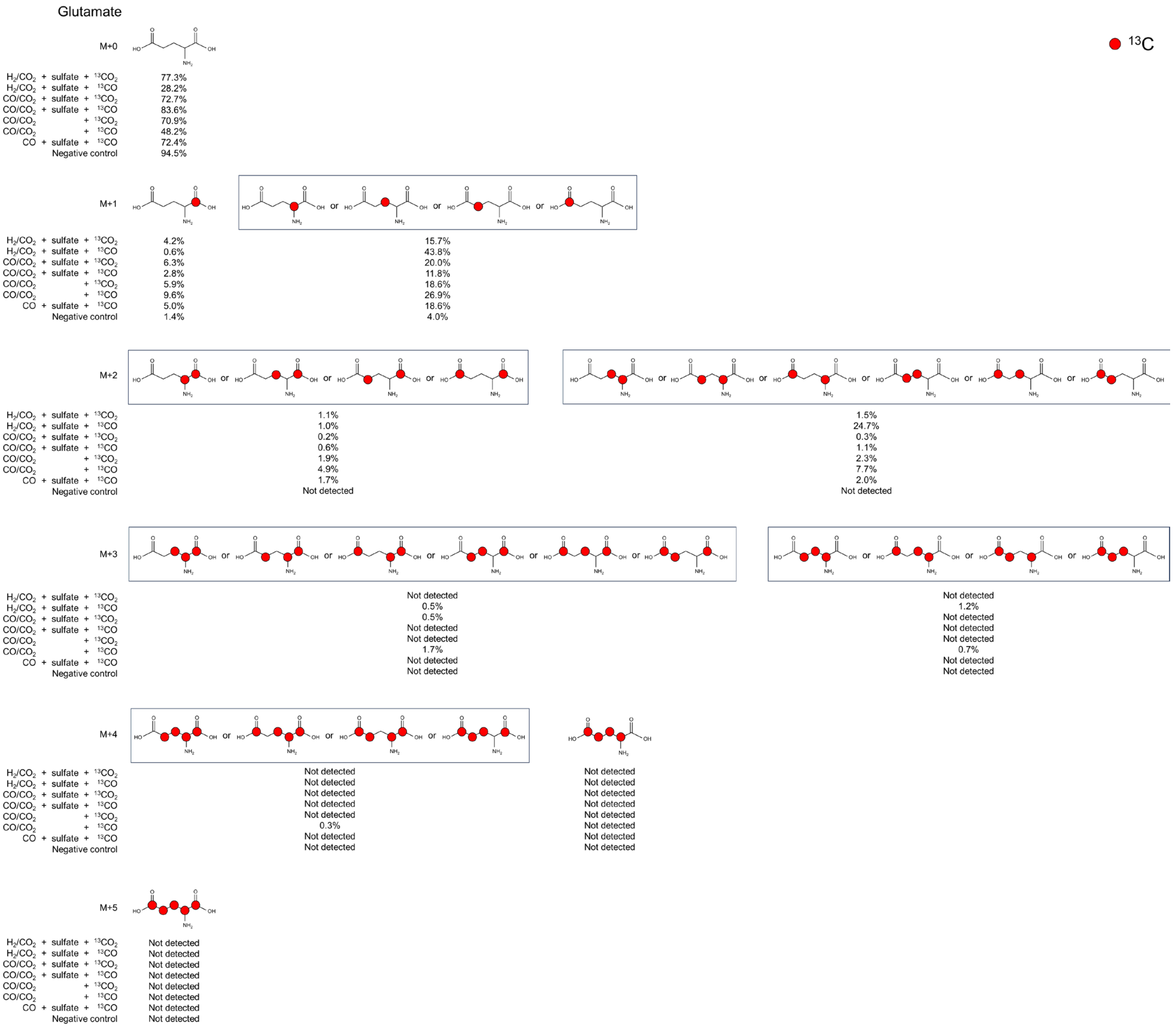


| | Isotopomer pattern | | Natural abundances of isotopes | *A. lithotrophicus* strain MCR | | | | | | | Negative control |
|---|---|---|---|---|---|---|---|---|---|---|---|
| | | | | $H_2/CO_2$ | | CO/$CO_2$ | | | | CO | |
| | No. of $^{13}C$ | | | $^{13}CO_2$ + sulfate | $^{13}CO$ + sulfate | $^{13}CO_2$ + sulfate | $^{13}CO$ + sulfate | $^{13}CO_2$ | $^{13}CO$ | $^{13}CO$ + sulfate | |
| Amino acid | C-1 | Others | Abundance (%) | Abundance (%) | Abundance (%) | Abundance (%) | Abundance (%) | Abundance (%) | Abundance (%) | Abundance (%) | Abundance (%) |
| $^{13}C_0$ Glu | 0 | 0 | 94.6 | 77.3 | 28.2 | 72.7 | 83.6 | 70.9 | 48.2 | 72.4 | 94.5 |
| $^{13}C_1$ Glu | 1 | 0 | 1.1 | 4.2 | 0.6 | 6.3 | 2.8 | 5.9 | 9.6 | 5.0 | 1.4 |
| | 0 | 1 | 4.2 | 15.7 | 43.8 | 20.0 | 11.8 | 18.6 | 26.9 | 18.6 | 4.0 |
| $^{13}C_2$ Glu | 1 | 1 | 0.0 | 1.1 | 1.0 | 0.2 | 0.6 | 1.9 | 4.9 | 1.7 | N.D. |
| | 0 | 2 | 0.1 | 1.5 | 24.7 | 0.3 | 1.1 | 2.3 | 7.7 | 2.0 | N.D. |
| $^{13}C_3$ Glu | 1 | 2 | 0.0 | N.D. | 0.5 | 0.5 | N.D. | N.D. | 1.7 | N.D. | N.D. |
| | 0 | 3 | 0.0 | N.D. | 1.2 | N.D. | N.D. | N.D. | 0.7 | N.D. | N.D. |
| $^{13}C_4$ Glu | 1 | 3 | 0.0 | N.D. | N.D. | N.D. | N.D. | N.D. | 0.3 | N.D. | N.D. |
| | 0 | 4 | 0.0 | N.D. | N.D. | N.D. | N.D. | N.D. | N.D. | N.D. | N.D. |
| $^{13}C_5$ Glu | 1 | 4 | 0.0 | N.D. | N.D. | N.D. | N.D. | N.D. | N.D. | N.D. | N.D. |

N.D., not detected

**Fig. S11 The relative abundance of isotopomer pattern in Glu from *Archaeoglobus* sp. strain MCR**

The mass fractions for M+0, M+1, M+2, M+3, M+4, and M+5 represent fragments containing 0 to 5 $^{13}C$-labeled carbons, respectively. Red circles show $^{13}C$-labeled carbon atoms in the carbon skeleton.
